# Natural variation in *Prdm9* affecting hybrid sterility phenotypes

**DOI:** 10.1101/2023.01.17.524418

**Authors:** Khawla FN AbuAlia, Elena Damm, Kristian K Ullrich, Amisa Mukaj, Emil Parvanov, Jiri Forejt, Linda Odenthal-Hesse

## Abstract

PRDM9-mediated reproductive isolation was first described in offspring of *Mus musculus musculus* strain PWD/Ph and *Mus musculus domesticus* strain C57BL/6J. Male F_1_-hybrids do not complete chromosome synapsis and arrest meiosis at Prophase I. Currently, all data supports an oligogenic control of hybrid sterility based on incompatibilities between PRDM9 and hybrid-sterility locus *Hstx2* in *Mus musculus* hybrids. Erosion of PRDM9 binding sites was proposed to result in asymmetric binding on diverged homologs of intersubspecific F_1_ hybrids. Numerous alleles of *Prdm9* have been characterized for different subspecies of *Mus musculus*, but only a few were analyzed for their impact on hybrid sterility. We analyzed *Prdm9* diversity in natural wild mouse populations from Europe, Asia, and the Middle East and identified several novel *Prdm9* alleles. We established that a single *Prdm9* allele is associated with *t*-haplotype Chromosome 17 in all three subspecies of *Mus musculus* and characterized the phylogenetic relationships of novel *Prdm9* alleles with established sterility alleles. Novel wild *Prdm9* alleles produced F_1_-hybrid male offspring that were either fertile or showed *Prdm9*-dependent reduction of fertility and high levels of asynapsis. Fertility or sterility phenotypes segregated purely with the *Prdm9* genotype, although the *Mus musculus musculus* background varied. Our data substantiate that hybrid sterility is under oligogenic control with *Prdm9* as the leading player but is consistent with a nonbinary regulation of hybrid sterility and gradual fertility decline when homologs diverge.

## Introduction

Hybrid sterility is an evolutionary concept of reproductive isolation in which hybrid zygotes develop into healthy adults who fail to produce functional gametes and are thus sterile. In the “Bateson-Dobzhansky-Muller model of incompatibilities”, hybrid sterility occurs when two evolved genes are incompatible upon interacting within an individual (Bateson 1909; Dobzhansky 1936; Muller 1942). The first identified hybrid sterility locus in mammals was Hybrid sterility 1 (*Hst1*) on chromosome 17 (Forejt and Ivanyi 1974). At the *Hst1* locus, the *Prdm9* gene is responsible for the observed hybrid sterility and codes for PR domain-containing protein 9 (PRDM9) (Mihola *et al*. 2009). The PRDM9 protein is expressed in testicular tissue during the early phases of meiotic Prophase I when recombination is initiated. It has three conserved domains, an N-terminal KRAB-domain that promotes protein-protein binding (Imai *et al*. 2017; Parvanov *et al*. 2017; Wang *et al*. 2021), an NLS/SSXRD repression domain with nuclear localization signal, and a central PR/SET domain providing methyltransferase activity (Powers *et al*. 2016). The C-terminal domain is highly polymorphic and comprises an array of C_2_H_2_-type zinc fingers (ZNFs) that differ between PRDM9 variants in both type and number (Baudat *et al*. 2010; Berg *et al*. 2010; Parvanov *et al*. 2010; Berg *et al*. 2011; Baudat *et al*. 2013; Buard *et al*. 2014; Kono *et al*. 2014). Variation between the ZNFs is most pronounced in the amino acids at positions −1, 3, and 6 of the ZNF α-Helix, responsible for recognizing specific DNA target motifs (Oliver *et al*. 2009; Baudat *et al*. 2010). Amino acid substitutions in positions −5, −2, and 1 of the α-Helix are rarely also seen but not predicted to affect protein-DNA binding affinity (Parvanov *et al*. 2010; Kono *et al*. 2014). Upon interaction with its specific DNA motif, the PR/SET domain of PRDM9 tri-methylates the adjacent nucleosomes on histone-3 by lysine-4 (H3K4) and lysine-36 (H3K36) (Hayashi *et al*. 2005; Wu *et al*. 2013; Eram *et al*. 2014), inducing a cascade of events that initiate recombination, as reviewed in (Damm and Odenthal-hesse 2022).

PRDM9-mediated reproductive isolation was discovered in male F_1_-hybrid offspring of *Mus musculus musculus* (MUS) and *Mus musculus domesticus* (DOM) strains, differing in *Prdm9* alleles (Forejt and Ivanyi 1974; Mihola *et al*. 2009). The PWD/Ph (hereafter PWD) strain possesses the *msc1* allele, and the C57BL6/J (hereafter B6) strain has the *dom2* allele. F_1_ male hybrids of PWDxB6 crosses do not complete chromosome synapsis, arresting at meiotic prophase I (Mihola *et al*. 2009; DZUR-GEJDOSOVA *et al*. 2012; Flachs *et al*. 2012). Defective pairing and high levels of chromosomal asynapsis are observed in hybrids with a large number of asymmetric sets of DSBs (Davies *et al*. 2016; Forejt 2016; Zelazowski and Cole 2016). When ineffective DSB repair prevents intersubspecific hybrids from forming gametes, a barrier of postzygotic isolation is created (Mihola *et al*. 2009; Davies *et al*. 2016). It was hypothesized that the molecular mechanism of PRDM9 action is related to the evolutionary divergence of homologous genomic sequences in *DOM* and *MUS* subspecies (Davies *et al*. 2016) and, more specifically, to the phenomenon of historical erosion of genomic binding sites of PRDM9 ZNF domains (Baker *et al*. 2015). Nucleotide polymorphisms within the PRDM9 target sequence can alter the binding affinity of PRDM9. When PRDM9 binding occurs more efficiently on a given DNA sequence motif, this motif is then consequently cut by DSB formation. However, as the uncut strand provides the template for repair, the less-efficient motif is repeatedly over-transmitted in heterozygous individuals and preferentially transmitted to the next generation. Therefore, polymorphisms that reduce the binding affinity of a specific PRDM9 variant become enriched within populations over time, resulting in the attenuation of hotspots (Boulton *et al*. 1997). Direct evidence of hotspot erosion has been observed in human and mouse hotspots (Jeffreys and Neumann 2005; Berg *et al*. 2011; Cole *et al*. 2014; ODENTHAL-HESSE *et al*. 2014). Indeed, in MUSxDOM hybrids, initiation asymmetry is mainly driven by functional *dom2* binding sites found on the PWD genome that are eroded on the B6 genome and vice versa for *msc1* sites (Davies *et al*. 2016). Further support for this hypothesis comes from the observation that fertility can be restored when the B6 PRDM9 zinc-finger array is replaced with that of the human variant B, making symmetric recombination hotspots predominant (Davies *et al*. 2016). This points to an oligogenic control, with PRDM9 as the main factor. This is also supported by findings relating the degree of chromosome asynapsis of PWD x B6 hybrids to meiotic arrest and the number of expected symmetric DSB hotspots per chromosome (Gregorova *et al*. 2018). Here, asynapsis was shown to operate in-*cis*, depending on increased heterozygosity of homologs from evolutionarily diverged subspecies. The introgression of at least 27 Mbs of sequence homology belonging to the same subspecies (consubspecific) fully restored the synapsis of a given autosomal pair (Gregorova *et al*. 2018). *Prdm9* is also a major hybrid sterility gene in MUS x CAS hybrids, where the synapsis rate was proportional to the level of nonrecombining MUS genetic background (Valiskova *et al*. 2022).

*Prdm9* effect on fertility of intersubspecific hybrids is modified by the X-linked locus *Hstx2*, located in a 2.7 Mb region on the proximal part of the X chromosome. *Hstx2* is structurally distinct between PWD and B6 mice and causes complete hybrid sterility only when the maternal *HstX2*^*PWD*^ is active (Bhattacharyya *et al*. 2014; Balcova *et al*. 2016). Here the interaction between *Prdm9* and the MUS *Hstx2* allele from the PWD strain enhances asynapsis of homologous chromosomes and eventually meiotic arrest (Bhattacharyya *et al*. 2013; Bhattacharyya *et al*. 2014; Balcova *et al*. 2016). In contrast, male F_1_ hybrids of the reciprocal cross (B6 X PWD) retain a low level of fertility. The *Hstx2* locus behaves like a recombination cold spot in crosses (Balcova *et al*. 2016), has reduced *Prdm9*-mediated H3K4me3 recombination initiation sites, and lacks DNA DSB hotspots decorated by DMC1 (Lustyk *et al*. 2019).

Hybrid sterility also occurs outside of laboratory models, as *Prdm9* alleles found among MUS and DOM wild-derived strains (Pialek *et al*. 2008) showed fertility disruption in about one-third of the intersubspecific male hybrids (Mukaj *et al*. 2020). Here, reduced sperm counts and low paired testes weights were associated with high asynapsis rates of homologous chromosomes in Meiosis I and early meiotic arrest in mice with *Prdm9* alleles that were closely related to previously identified hybrid sterility alleles (Mukaj *et al*. 2020). Replacing *dom2* for the “humanized” *Prdm9* transgene *Prdm9*^*tm1*.*1*^*(PRDM9)*^*Wthg*^ from *(Davies et al. 2016)* completely restored fertility in these mouse hybrids of wild-derived inbred strains, corroborating the role of PRDM9 as the leading player (Mukaj *et al*. 2020). In this study, the same *Prdm9* allele could act as hybrid sterility in one strain but not another, and the influence of the genomic background on this phenomenon remains unknown. In the natural hybrid zone, a relatively young zone of secondary contact, only as few as one-third of all house mouse males displayed fertility traits below the range of the pure subspecies (Turner and Harr 2014). Five *Prdm9*-dependent quantitative trait loci have been identified in intersubspecific (MUS x CAS) hybrids, segregating on DOM background (Valiskova *et al*. 2022). Several interchangeable autosomal loci have been proposed that may suffice to activate the Dobzhansky-Muller incompatibility in wild mouse hybrids (DZUR-GEJDOSOVA *et al*. 2012; Turner and Harr 2014), but they may be independent of *Prdm9*. In wild mice, *t* haplotypes, 30 Mb of a naturally occurring chromosome 17 haplotype (Silver 1985) also strongly influence male fertility in MUS and DOM mice. The t-haplotype, which has introgressed into *Mus musculus* subspecies from an unidentified *Mus* antecedent over one million years ago (Hammer and Silver 1993), encompasses the *Prdm9* locus (Trachtulec *et al*. 2008), and *Prdm9* located within the *t*-haplotype region, displays one of the most diverse alleles identified in the *Mus musculus* subspecies to date (Kono *et al*. 2014). However, it is not known whether *Prdm9* contributes to the observed reduction in fertility associated with *t*-haplotypes. Males heterozygous for the *t*-haplotype pass it on to more than half of their offspring, with some variants presenting transmission rates over 90%, while females transmit the *t*-haplotype within the expected Mendelian ratio (Lyon 2003). Despite a strong drive, *t*-haplotypes are only present in 10-40% of all subspecies of wild house mice, presumably because they also include genes causing male infertility and embryonic lethality (Olds-clarke 1997; Planchart *et al*. 2000; Schimenti *et al*. 2005; Kelemen and Vicoso 2018). Several distorter loci of the t-haplotype act in trans to impair motility in wildtype spermatozoa by over-activating a signaling pathway that controls sperm motility kinase (SMOK), resulting in abnormal flagellar movements and loss of sperm motility (Herrmann *et al*. 1999). To prevent the distorter locus from affecting t-haplotype-bearing sperm, a responder locus composed of a fusion gene of a SMOK member and the 3’-UTR of the triple ribosomal s6 gene conferring partial cis-resistance to overactivation by the distorters, thus restoring sperm motility (Herrmann and Bauer 2012)

As historical erosion of hotspot motifs has been postulated as the source of *Prdm9*-mediated sterility, we inquired into the evolutionary age of alleles using phylogenetic reconstructions of the *Prdm9* hypervariable minisatellite, coding for the ZNF array responsible for DNA binding affinity of the PRDM9 protein. To investigate whether *Prdm9*-mediated reproductive isolation operated in wild mice, we tested alleles from natural wild mouse populations from Europe, Asia, and the Middle East, including *Prdm9*-alleles on introgressed t-haplotypes, for whether they affected fertility in intersubspecific hybrids. Our results are consistent with the low levels of hybrid sterility phenotypes seen in wild mice and provide the first evidence that full hybrid sterility segregated by *Prdm9* genotype even on wild genomic backgrounds. Together with previous findings (Mukaj *et al*. 2020; Valiskova *et al*. 2022), our data substantiate that intrasubspecific variation in the genomic background does not explain the low incidence of hybrid sterility phenotypes seen in wild mice.

## Material and Methods

### Mice

All work involving experimental mice was performed according to approved animal protocols and institutional guidelines of the Max Planck Society and with permits obtained from the local veterinary office ‘Veterinäramt Kreis Plön’ (permit number: 1401-144/PLÖ-004697). Mice, including strains of PWD/Ph strain, C57/Bl6 strain with transgene *Prdm9*^*tm1*.*1*^*(PRDM9)*^*Wthg*^ strain and consomic C57BL/6J-Chr X.1s^PWD/Ph^/ForeJ mice, as well as several wild mice populations were all maintained in the mouse facilities of the Max Planck Institute for Evolutionary Biology in Plön, following FELASA guidelines and German animal welfare law. We analyzed three outcrossed populations of DOM mice, first the French Massif-Central (MCF) population, founded in December 2005 with a starting population size of sixteen breeding pairs, with additional wild-caught animals introduced into the breeding population at the beginning of April 2010. These mice were in generation sixteen at the start of this experiment, nine generations since crossing in with the second set of new wild-caught animals. The German Cologne-Bonn (CBG) population was founded in August 2006 with ten breeding pairs and maintained as an outcross for fourteen generations at the start of this experiment. In November 2012, new wild-caught breeding pairs were crossed in. The Iranian population from Ahvaz (AHI) was started in December 2006, with six founding breeding pairs. It has been maintained in an outcross for fifteen generations at the start of this experiment, after two rounds of reduction due to inbreeding depression. Seventeen breeding pairs of mice originally trapped in Almaty, Kazakhstan (AKH) in December 2008, founded the *Mus musculus musculus* population, which had been maintained for thirteen generations at the beginning of this experiment.

### Organ withdrawal

Organ withdrawal after euthanasia is not legally considered an animal experiment according to §4 of the German Animal Welfare Act. It therefore does not need to be approved by the competent authority (Ministerium für Landwirtschaft, ländliche Räume, Europa und Verbraucherschutz). F_1_ hybrid males were euthanized after they were first rendered unconscious by the deliberate introduction of a specific CO_2_/O_2_ mixture ratio, then sacrificed using CO_2_ euthanasia followed by cervical dislocation. To reduce loose hair contaminating the organs during the dissection of the animal, their coat was sprayed with 75% EtOH before organ withdrawal. Spleen, a liver lobe, and both testes were extracted, and epididymides were removed. One epididymis was placed in 1 ml of cold phosphate-buffered saline, and all other organs were immediately snap-frozen in liquid nitrogen and stored at −70° C.

### Fertility phenotyping

We collected three fertility parameters, body weight (BW) and paired testes weight (TW), and spermatozoa released from epididymal tissues, counted in Million/ml (SC). One epididymis, including caput, corpus, and cauda, was repeatedly cut in 1 ml of cold phosphate-buffered saline to release spermatozoa. The tube was vigorously shaken for 2 minutes, and spermatozoa in the solution were diluted to 1:40 in PBS. We counted 10 µl of diluted spermatozoa in a Bürker chamber (0,1 mm chamber height), where two replicates of 25 squares were counted. In cases when only a few (<10) spermatozoa were found, additional dilutions were prepared and counted. To approximate spermatozoa released from a pair of epididymides, we added the two replicated 25 squares counts (A_25 +_ B_25_) from spermatozoa released from a single epididymis. The epidydimal spermatozoa count released in 1 ml PBS was then calculated by taking the paired counts, the volume of 25 squares (V_25_=0,02*0,02*0,01*25=0,0001 cm^3^), and the dilution factor into account.

### Spreading and Immunofluorescence analyses of spermatocytes

Spermatocyte nuclei were spread for immunohistochemistry as described in (Anderson *et al*. 1999), with the following modifications. Firstly, a single-cell suspension of spermatogenic cells from the whole testis was prepared in 0.1 M sucrose solution. The sucrose-cell slurry, with added protease inhibitors (Roche 11836153001), was then dropped onto paraformaldehyde-treated glass slides. Glass slides were kept in a humidifying chamber for 3 hours at 4 °C to allow cells to spread and fix. Slides were briefly washed in distilled water and transferred to pure PBS before blocking in PBS with 5-vol% goat serum. Primary antibodies HORMAD2 (a gift from Attila Toth, rabbit polyclonal antibody 1:700), SYCP3 (mouse monoclonal antibody, Santa Cruz, #74569, 1:50), yH2AX (ab2893. 1:1000), and CEN (autoimmune serum, AB-Incorporated, 15-235) were used for immunolabelling. Secondary antibodies goat anti-Mouse IgG-AlexaFluor568 (MolecularProbes, A-11031), goat anti-Rabbit IgG-AlexaFluor647 (MolecularProbes, A-21245), goat anti-Human IgG-AlexaFluor647 (MolecularProbes, A-21445), goat anti-Rabbit IgG-AlexaFluor488 (MolecularProbes, A-11034) were used at 1:500 concentration at room temperature for one hour. A Nikon Eclipse 400 microscope with a motorized stage control was used for image acquisition with a Plan Fluor objective, 60x (MRH00601). Images were captured with a DS-QiMc monochrome CCD camera and the NIS-Elements program (from Nikon). Image J software was used to process the images.

### PRDM9 Genotyping

We used ear clips taken at weaning to identify *Prdm9* allelic variation in the wild mouse populations. All F_1_ and F_2_ hybrid offspring used in the experiments were instead genotyped from the counted sperm sample taken from one of the epididymides. Furthermore, we confirmed initial parental PRDM9 genotyping after successful mating (> 5 male offspring) by sacrificing all F_0_ males. All genotyping was done on individual mouse IDs, but in such a way that the experimenter was blind to the matching fertility phenotypes.

### DNA extraction

DNA was extracted from ear clips or whole ears using salt extraction. Briefly, cells were lysed in SSC/0.2 % SDS, and proteins were digested using Proteinase K (20 mg/µl), incubating at 55 °C overnight. We salted out the DNA using 4.5 M NaCl solution, followed by two consecutive rounds of Chloroform extraction. The DNA was then Ethanol precipitated and washed twice with 70 % ethanol, and the pellet was then dried at room temperature and finally dissolved in 30 µl Tris-EDTA pH 8.0. The DNA samples were stored at 4 °C for short-term storage and −70 °C long-term. The slurry of isolated spermatozoa with epididymal tissues was processed similarly; however, to lyse sperm heads and remove Protamines, we increased the SDS concentration to 1 % and added not only Proteinase K (20 mg/µl) but also TCEP (Thermo Scientific 77720, 0.5 M) to a final concentration of 0.01 µM. This extraction method produces a mixture of DNA extracted from somatic and sperm cells.

### Amplification of the minisatellite coding for the zinc-finger array of PRDM9

The ZNF arrays of each mouse were PCR amplified similarly as in (Buard *et al*. 2014) on 10-30 ng of genomic DNA in 12 µl reactions of the PCR buffer “AJJ” from (Jeffreys *et al*. 1990) using a two-polymerase system with Thermo Taq-Polymerase (EP0405) and Stratagene Pfu Polymerase (600159) to ensure d high-fidelity PCR. When offspring are heterozygous for two alleles of different lengths (in most cases), we separated heterozygous bands after gel electrophoresis on Low Melting agarose (Thermo Fischer #R0801) by excising the bands and eluting the DNA using Agarase (Thermo Fischer #EO0461). If two heterozygous bands were apparent, excised and eluted product was immediately used in sequencing reactions after estimating the amount of DNA from the gel. If only one band was evident, alleles were not separated by size. Therefore, the purified PCR products were cloned using TOPO TA Cloning Kit for Sequencing (Life Technologies no. 450030), following the manufacturers’ specifications before sequencing. We analyzed at least eight clones per sample.

### Sequencing

Sequencing reactions of either eluted PCR product or picked clones were set up using BigDye 3.0, according to the manufacturer’s protocol, then purified using X-terminator, and finally sequenced using 3130x/ Genetic Analyser. Only PRDM9 variants with less than 12 ZNFs (sequence lengths of >1200 bp) could be sequenced up to their ends in both directions. We assembled the forward and reverse sequences based on the estimates of fragment sizes from PCR products on gels to accurately assemble the coding minisatellite using Geneious Software (Version 10.2-11). After sequencing and alignment, assembled minisatellites were conceptually translated into the amino acid sequence of the ZNF domain, and HMMER scores were computed using a Polynomial SVM (Persikov and Singh 2014).

### Phylogenetic Analyses

The phylogeny on all alleles tested for hybrid sterility phenotypes in (Mukaj *et al*. 2020) and this publication was computed using the R package “repeatR” from https://mpievolbio-it.pages.gwdg.de/repeatr/ (Damm *et al*. 2022). Briefly, minisatellite-like repeats within the gene are identified, extracted, and filtered for incomplete sequences before matrices based on minimum edit distance (Hamming) were computed using weighting costs w_mut_=1, w_indel_=3.5 and w_slippage_=1.75 as given in (Vara *et al*. 2019). These Minimum edit distances represent a metric on the set of changes between *Prdm9* minisatellite repeat units of 84 bp in lengths. As such, can be used as a measure of genetic distance. We computed two distance matrices for each type of repeat, as in (Damm *et al*. 2022). The first distance matrix included all nucleotides, while the second matrix excluded nucleotides known to be under positive selection, which is those coding for the hypervariable amino acids responsible for DNA binding specificity (−1, +3, +6). Two phylogenetic reconstructions of the *Prdm9* hypervariable region were then computed separately from both matrices, using a neighbor-joining approach with the “bionj” function of the R package ape [79], and rooted on the” humanized” *Prdm9* allele from (Davies *et al*. 2016).

### Genotyping for Chr17 t-haplotype and X-chromosomal haplotypes near *Hstx2*

The presence of the t-haplotype was tested using markers Tcp1 and Hpa-4ps (Planchart *et al*. 2000), and X-chromosomal haplotypes across the refined *Hstx2* interval were tested using primers from (Lustyk *et al*. 2019), **Table S1**. Each forward primer was labeled with either HEX or FAM and amplified using the ABI Multiplex Kit according to the manufacturer’s protocol. Fragment lengths were then analyzed by capillary electrophoresis using a 3730 DNA Analyser. Allele sizes were scored and binned using the Microsatellite plugin in Geneious v.10.2.

### Statistical Analyses

The majority of graphs, calculations, and statistical analyses were performed using GraphPad Prism software version 9.4.1 for Mac (GraphPad Software, San Diego, CA, USA). Statistical tests are stated in the text as well as in the Figure legends. Briefly, pairwise comparisons were performed using unpaired t-tests with Welch correction, and as we did not assume equal sample variances *a priori*, these were compared using F-tests. Similarly, multiple comparison tests of fertility parameters were also first evaluated for differences in sample variances using Brown-Forsythe ANOVA tests. If significant differences between means were observed, we performed Welch’s ANOVA with Dunnett’s T3 multiple comparisons test. When there was no indication of unequal sample variance, we performed ordinary one-way ANOVA instead, which we evaluated with Bonferroni multiple comparisons tests. Asynapsis data were compared between genotypes also using unpaired t-tests with Welch correction, and linear correlation was assessed using the Pearson correlation coefficient (r). We tested for transmission ratio distortion using the binomial probability calculator on the VassarStats: Website for Statistical Computation (http://vassarstats.net). In general, only significant values are shown on graphs with *P*<0.0332 (*), *P*<0.0021(**), *P*<0.0002(***), and *P*<0.0001(****).

## Results and Discussion

Previous analyses had identified several alleles of *Prdm9* that induced hybrid sterility in European mice *(Mukaj et al. 2020; Forejt et al. 2021)*. We screened additional European wild mice further away from the hybrid zone as well as mice from Asia and the middle-east for novel *Prdm9* alleles. Mice were initially caught by *(HARR et al. 2016)*, and all populations have been housed and maintained as an outcrossed population over many generations ahead of this study (for details, see Materials and Methods). MUS were originally trapped in Almaty, Kazakhstan (43°16’N, 76°53’E), and DOM in three different locations; the city of Ahvaz, Iran (31°19’ N, 48°42’ E), the Massif-Central area in France (45°32’N, 2°49’E) and the Cologne-Bonn area in Germany (50°52’N, 7°8’E). The distribution of original trapping locations of all mice tested for hybrid sterility phenotypes, in this study and in (Mukaj et al. 2020) are shown in **Figure S1**.

We screened all populations for *Prdm9* alleles by sequencing exon12 of *Prdm9* containing the minisatellite coding for the C_2_H_2_ zinc-finger domain of PRDM9 as described in (Buard et al. 2014; Kono et al. 2014). The amino-acid variation between ZNF domains was determined by conceptually translating the nucleotide sequence of satellite repeats into the amino-acid sequence of individual zinc fingers (**Figure S2** and **Figure S3**). We defined the ZNF variation based on the amino acids present at positions −1, +3, and +6 of the alpha-helix, representing the DNA contact residues (**Figure S4A**). Variants with amino acid changes outside of the alpha-helix are not predicted to have altered DNA binding specificity. Some repeats possess a Tryptophan (W) residue in position −5. Tryptophan’s nonpolar, aromatic and neutral chemical properties which differ from the Arginine (R) typically found in this position, which is polar and strongly basic. Secondly, there are two types of last repeats; the rarer one possesses Arginine (R) in position 13, while the more common type contains aliphatic and nonpolar Glycine (G) (**Figure S2**). Glycine appears to be the ancestral allele, as it is located in this position also in human *Prdm9* allele B (**Figure S3**), able to rescue sterility when genetically engineered into mice (Davies *et al*. 2016).

Based on variation in nucleotide repeats, as well as the composition, order, and number of repeats, we have identified eight full-length *Prdm9* alleles in MUS mice from Kazakhstan and six in DOM from France, Germany, and Iran (in **Figure S4B**). We named these *Prdm9* alleles according to the International Committee of Standardized Genetic Nomenclature for mice (MGI) (in **Figure 1** and **Figure S4B**), registered them at JAX, and submitted their sequences to GenBank under Accession numbers (OQ055171-OQ055188).

**Figure 1.**
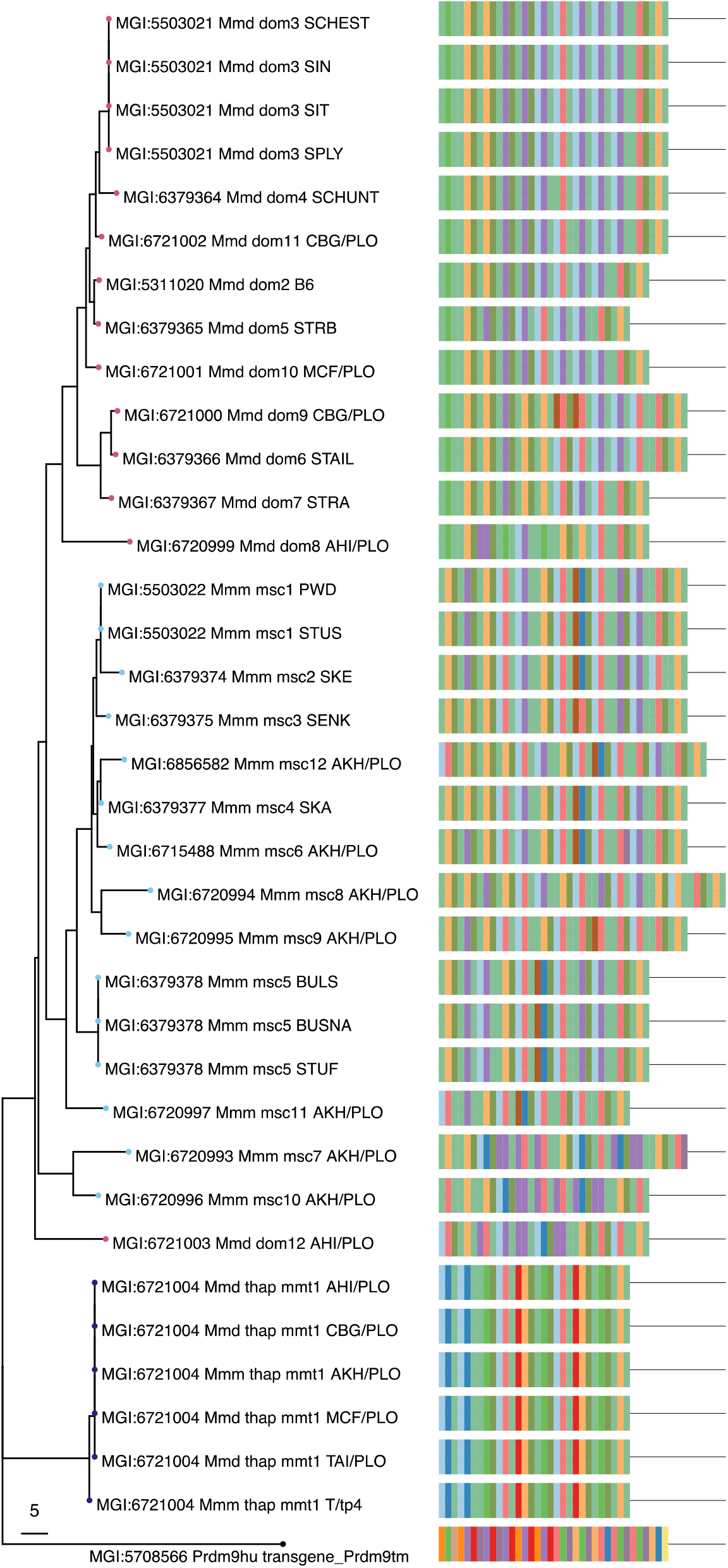
Diversity and distribution of PRDM9 variants tested for hybrid sterility. Allelic diversity of *Prdm9* in mice in this study and, as well as the PRDM9^Hu^ “humanized” from (Davies *et al*. 2016) **(tree) neighbor-joining** tree based on Hamming-Distances computed on the full-length nucleotide repeat sequences coding for C_2_H_2_ ZNFs of PRDM9 using repeatR (Damm *et al*. 2022). **(#ZNF)** the length of the ZNF array **(−1, +3, +6) color-coded** PRDM9 zinc finger domains are characterized by the type of C_2_H_2_ zinc fingers, listing the variable amino acids at DNA binding positions −1, +3, +6 determining DNA binding specificity (as shown in **Figure S2 Error! Reference source not found**.).

We found a peculiar *Prdm9* allele in both MUS and DOM subspecies in all four original trapping locations in mice who also possessed t-haplotypes. In this allele, nine nucleotides are deleted in the first ZNF of the array, removing three amino acids in the translated amino-acid sequence, including one of the zinc-binding Cysteine ligands (**Figure S2**). In addition, distinct amino acids are seen in positions −1, +3, and +6, not present in any other PRDM9 variants, such as TDK and ASQ, with additional differences between the two types of ASQ ZNFs in positions −8 and +5. Similarly, the ANQ ZNF found in mice with *t*-haplotypes differs from the ANQ found in mice without *t*-haplotypes at position −2. This allele had also been previously found in mice with t-haplotypes (Kono *et al*. 2014). We, therefore, tested additional mice with t-haplotypes for *Prdm9*, including *Mus musculus castaneous* (CAS) from Taiwan and mouse strain T/tp4 (Forejt et al. 1988), confirming that both also possessed the same *Prdm9* allele. Given that a single *Prdm9* allele was found in all mice with t-haplotypes, regardless of subspecies, we considered it to be an intraspecies *Mus musculus* allele and named it *Mus musculus* t-haplotype *Prdm9* allele 1 “*mmt1”*. Together with t-haplotypes, the *mmt1* allele was always present in a heterozygous state and occurred in all of our outcrossed populations at high frequencies. We found t-haplotypes in 50% of the MUS population from Almaty (Kazakhstan), in 88% of the DOM population from Ahvaz (Iran), in 90% of the mice from Cologne-Bonn (Germany), and in 100% of the mice from the Massif-central (France) population. As mice had been outcrossed over many generations before this study was initiated, these high population frequencies do not reflect t-haplotype frequencies in the wild.

All other alleles in this study were present only in one subspecies and only in a single population, but some had been identified previously. Allele *msc11* had been found in a MUS from Illmitz, Austria, where it was named *7mus1* (Buard *et al*. 2014) and closely resembled two alleles from the CAS subspecies. Firstly, the classical *Cst* allele possesses Serine at position −1 of the alpha-helix of the 8^th^ ZNF (Parvanov *et al*. 2010); in this position, *msc11* has a single amino-acid change to Asparagine. Similarly, a CAS trapped in Nowshahr, Iran, *Ca1* (Kono *et al*. 2014), differs only by a single amino-acid substitution. In position 6 of the alpha-helix of the 6^th^ ZNF, *Ca1* possesses Glutamine, while *msc11* possesses Lysine instead. Allele *msc9* was found in MUS strains CHD and BLG2, where it was named *Ma12* (Kono *et al*. 2014). Allele *msc6*, which we found in mice from Kazakhstan, has also previously been identified as Ma8 in MUS from Grozny, Russia (Kono *et al*. 2014) and appears closely related to the *msc1* allele from PWD and STUS/JPia strains, differing only in two single-nucleotide polymorphisms that lead to differences in amino-acids of ZNF11 (as shown in **Figure S4B**). Alleles msc7, *msc8, msc10*, and *msc12* are novel but bear considerable similarities with other previously identified MUS alleles. The *msc12* allele appears closely related to the *msc4* allele from the SKA/JPia strain, differing only in the addition of a proximal ZNF. Alleles msc7 and *msc10* appear similar to each other, MUS 27mus1 trapped in Bulgaria (Buard *et al*. 2014), and *Cc4* rapped in Grozny, Russia (Kono *et al*. 2014).

The alleles found in our DOM mice share similarities to previously identified DOM alleles. The *dom8* allele, which we found in Ahvaz, Iran, has been identified before in the DOT strain, originally from Tahiti (French Polynesia), where it was named 16dom1 (Buard *et al*. 2014). The remaining DOM alleles are all novel. Allele *dom9* found in the Cologne-Bonn area of Germany is, however, highly similar to the *dom6* allele found in the STAIL/JPia strain, initially trapped in Schweben, Germany. The *dom10* found in Massif-Central is similar to the *dom3* allele found in strains SPLY/JPia trapped in Bavaria, Germany, and the SIN/JPia strain trapped in the Orkney Islands, UK, lacking only the 7^th^ ZNF, compared. Similarly, the *dom11* allele, which we found in Cologne-Bonn, Germany, is similar to the *dom4* allele from the SCHUNT/JPia strain caught in Hesse, Germany, and even more similar to *dom3*, differing exclusively at a single amino-acid position −1 of the 9^th^ ZNF, where it possesses Glutamine instead of the Alanine, more common in this position (as shown in **Figure S4B**). The *dom12* allele identified in mice from Ahvaz, Iran, is the most unusual of the DOM alleles; it is identical to the CAS allele *Ce1* found in China (Kono *et al*. 2014) and shares high similarities to MUS allele *Cc4* found in Grozny, Russia.

### Phylogenetic analyses of *Prdm9* alleles in mice

It was proposed that erosion of PRDM9 binding sites results in asymmetric initiation when coupled with evolutionary divergent homologous genomic sequences of DOM and MUS subspecies in intersubspecific hybrids. Suppose the sterility of *Prdm9* is indeed driven by historical erosion of binding sites. In that case, alleles with a lower divergence from the last common ancestor are likely older alleles and, as such, may have suffered larger levels of historical erosion. To enquire into the evolutionary history of *Prdm9* alleles, we analyzed the phylogenetic relationship of alleles present in our wild mice populations, other alleles for which *Prdm9*-mediated hybrid sterility had been studied (Parvanov *et al*. 2010; Mukaj *et al*. 2020), and as outgroup added the humanized *Prdm9* “B-allele” (Davies *et al*. 2016). As handling sequence repeats is challenging for multiple-sequence alignment algorithms, particularly when the number of repeat units differs, the allelic divergence of minisatellite sequences cannot be assessed by standard assembly programs. In addition, genetic distance models (i.e., Tamura-Nei) do not accurately reflect minisatellite evolution, driven by *de-novo* recombination between repeats (Jeffreys *et al*. 2013). To reflect *Prdm9* evolution more accurately, we, therefore, applied an algorithm that computes Hamming Distances between minisatellite repeat, which not only takes point mutations and small indels but also within-repeat-unit processes (w_mut_=1), as well as repeat-unit insertions and deletions (w_indel_=3.5) and even repeat-unit duplication and slippage (w_slippage_=1.75) into account (Vara *et al*. 2019; Damm *et al*. 2022). For a more conservative phylogenetic analysis, we only included nucleotide repeats that, when translated into amino acids, had a Hidden Markov Model (HMMER bit) score above 17.7, determined using (Persikov and Singh 2014). This removed the nucleotide repeat coding for the first zinc finger in the ZNF array, which we found to be conserved in all *Prdm9* alleles except the *mmt1* allele.

In the neighbor-joining tree of Hamming distances rooted on the “humanized” *Prdm9* allele, alleles cluster broadly according to mouse subspecies (as shown in **Figure 1**). However, not all alleles follow the MUS/DOM subspecies divide. The *mmt1 Prdm9* allele found in all mice with *t-*haplotypes formed a separate branch irrespective of subspecies and mouse origin, a pattern typical of introgression. The large degree of conservation of *Prdm9* on the t-haplotype stands in stark contrast to the rapid evolution of *Prdm9* alleles in mice. Particularly, given the large divergence of *Prdm9* alleles in all natural populations studied to date, repeated introgression of the same allele at multiple independent events appears less likely than a single ancestral introgression event of t-haplotypes into an antecedent of all *Mus musculus* subspecies. Furthermore, recombination suppression on t-haplotypes due to a series of naturally occurring inversion blocks, including one close to the *Prdm9* locus (Kelemen and Vicoso 2018), likely constrains *Prdm9* evolution, which is mainly driven by *de-novo* recombination events during meiosis, including crossover and gene-conversion-like events (Jeffreys *et al*. 2013).

Except for the introgressed *mmt1* allele and the *dom12* allele, found neighboring a branch of MUS alleles and displaying low divergence to the last common ancestor of MUS and DOM alleles, all other alleles are separated by subspecies origin (as seen in **Figure 1** and **Figure S5**). The first node separates the *dom8* allele from the Iranian population from all other DOM alleles clustering by subspecies. The next node then separates several alleles from mice originally trapped in Germany, including alleles *dom7* from STRA/JPia and *dom6* from STAIL/JPia strains, as well as *dom9*, newly identified in the Cologne/Bonn, Germany population. Further nodes first separate the *dom10* allele from France and then a branch with *dom5* from STRB/JPia and hybrid sterility allele *dom*2, with comparable divergence. Another node first separates the *dom11* allele, closely followed by the *dom4* allele, found in SCHUNT/JPia strain, and finally, a branch with *dom3* alleles. The *dom3* alleles are found in SPLY/JPia, SCHEST/JPia, SIT/JPia, and SIN/JPia strains but represent a hybrid sterility allele only in the latter three strains.

A single node leads exclusively to all tested MUS alleles. Newly identified *msc10* and *msc7* alleles, which share considerable similarities to CAS alleles in a previous study (Kono et al. 2014), branch off first, followed by a branch harboring the novel *msc11* allele, with both branches showing the least divergence to the common ancestor of MUS alleles (as seen in **Figure 1**). Two nodes further separate the MUS-specific alleles into branches with larger divergence. One node exclusively leads to the three strains, BULS/JPia, BUSNA/JPia, and STUF/JPia, which all possess the *msc5* allele. The second node connects all other MUS *Prdm9* alleles. Listed with ascending divergence are the *msc4* allele from the SKA/JPia strain and newly identified *msc12* alleles, as well as *msc6*, located on a neighboring branch. Further separated are the *msc3* allele from the SENK/Jpia strain and the hybrid sterility allele *msc2* from the SKE/Jpia strain, followed closely by the *msc1* hybrid sterility allele from PWD and STUS/JPia. According to our phylogenetic reconstruction, all previously identified hybrid sterility alleles are subspecies-specific alleles of large divergence from a common ancestor.

However, our first approach makes interpreting divergence times difficult as it is known that the nucleotides coding for amino acids at positions −1, +3, and +6 of the alpha-helix of PRDM9 are under strong positive selection (Oliver *et al*. 2009). And as loci under positive selection can influence divergence times, we calculated a second distance matrix of Hamming distances after removing the hypervariable amino acids at −1, +3, and +6 of the alpha-helix, which are responsible for DNA binding before computing a second phylogeny. Indeed, several alleles also possess substitutions outside of the nucleotide positions coding for hypervariable amino acids responsible for DNA binding, and the topology of the tree changes dramatically (when comparing **Figure 1** and **Figure S5**). For alleles that differ outside of the positively selected sites, larger divergence times are not driven by positive selection alone, and they have thus apparently been present for longer evolutionary timescales. While the *mmt1* allele remains separated, some subspecies-specific nodes disappear. The closely related alleles to the common ancestor of *Mus musculus Prdm9* alleles are *msc11, msc5*, and *msc10*, which are now neighboring DOM alleles, possibly placing their origin before a clear separation into MUS and DOM subspecies. Curiously, while the full-length sequence of the *msc7* allele had previously appeared most closely related to the *msc10* allele (**Figure 1)**, it is now found neighboring the *msc2* allele from SKE/JPia, within a tree of subspecies-specific alleles (**Figure S5**). In contrast, alleles *msc1, msc4*, and *msc9* differed only at hypervariable sites, as they are found on the same branch once loci under positive selection are removed. Yet, the *msc6* allele remains separated as its nucleotide sequence has substitutions outside the hypervariable sites, pointing to longer divergence times between alleles. Similarly, without positively selected hypervariable sites, *dom6* and *dom9* are now located on the same branch. The divergence between *Prdm9* alleles thus appears predominantly driven by positive selection on the hypervariable sites. In conclusion, the complementary phylogenic analyses support the accelerated evolution of the hypervariable binding sites of *Prdm9* and reiterate an evolutionary history in which *Mus musculus* originated in Asia and the middle east before dispersing across Europe. The phylogenetic analyses further support a scenario in which MUS and DOM have split recently and are still speciating but do not reveal clustering of hybrid sterility alleles.

### Testing *Prdm9* alleles for sterility phenotypes in intersubspecific hybrids

To inquire whether newly identified wild *Prdm9* alleles would induce hybrid sterility, replicated crosses of the laboratory models of hybrids sterility but exchanged the father with a wild mouse male of the same subspecies. To test DOM alleles, PWD females (MUS) were crossed with wild DOM males, emulating the PWD x B6 laboratory model. To test MUS alleles, we emulate the B6.DX1s x PWD laboratory model (Mukaj *et al*. 2020). Here, we crossed wild MUS males to C57BL/6J-Chr X.1s^PWD/Ph^/ForeJ females (abbreviated B6.DX1s), which is a consomic DOM of C57Bl6/J strain background that carries the *Hstx2* locus within a 69.6 Mb PWD sequence of the proximal end of the X-chromosome, essential for F_1_-hybrid sterility (Bhattacharyya *et al*. 2014; Balcova *et al*. 2016; Lustyk *et al*. 2019; Forejt *et al*. 2021).

All nine DOM sires and five of the sixteen MUS sires possessed *t*-haplotypes. The *t-*haplotype is a known meiotic driver, skewing transmission against wildtype Chr17 in the male germline. We, therefore, experienced a severe reduction of testable alleles on wildtype Chr17 in both MUS and DOM mice, fully explained by the over-transmission of the *Prdm9*^*mmt1*^ allele with the t-haplotype. Indeed, more than 87% of offspring from fathers with *t*-haplotypes inherited the *mmt1* allele, a significant deviation from Mendelian 50:50 transmission ratio (two-tailed binomial probability *P=* <0.000001, approximated via normal). In contrast, *Prdm9* alleles present on wildtype chromosomes were transmitted at a 50:50 ratio in mice without *t*-haplotypes. As an expected outcome of comparing inbred and outbred mice are differences in fertility parameters, we determined and compared fertility parameters of the source populations of wild mice, which we compared with sperm counts and testes weights of inbred mice (**Figure S6**). Wild mice from some of our wild DOM populations had comparable sperm counts, while testes weights were significantly higher than in B6 mice, and similarly, wild MUS had elevated testes weights compared to PWD. As wild mouse sires stemmed from outcrossed populations, they were almost always heterozygous for different *Prdm9* alleles. Several males of the same population, which we used as sires in different crosses, shared one of the *Prdm9* alleles. Consequently, offspring from crosses with different fathers could obtain the same *Prdm9* allelic combination. We, therefore, wanted to investigate whether different genetic backgrounds of the fathers affected fertility parameters and did so by comparing sperm counts and paired testes weights between F_1_ hybrid offspring with the same *Prdm9* allelic combination, inherited from different fathers coming from the same source population. We did not observe significant differences between sperm counts (**Figure S7A)**, except in some offspring that inherited *t*-haplotypes (**Figure S7B**). Paired testes weights, however, were often significantly different between offspring from different fathers, particularly when they also carried *t*-haplotypes (**Figure S7**). When we pooled fertility parameters of F_1_-hybrid males by *Prdm9* genotype, most were significantly elevated compared to control hybrid sterility crosses PWD x B6 and B6.DX1s x PWD, with two notable exceptions. Intersubspecific hybrids that inherited paternal *msc11* and *msc12* alleles had the lowest sperm counts and testis weights, and their fertility parameters did not differ significantly from sterile controls (**Figure 2)**. Given that all F_1_-hybrid males were either completely fertile or showed only a reduction of fertility, we wanted to test whether they would display chromosomal asynapsis, a hallmark characteristic of *Prdm9*-dependent hybrid sterility. We then performed immunofluorescence analyses of synaptonemal complexes on spermatocyte spreads of F_1_-hybrids that inherited alleles *msc10, msc11*, and *msc12*, and hybrids that inherited the *mmt1* allele in combination with both *DOM* and *MUS* t-haplotypes. To quantify asynapsis, we scored each HORMAD2 stained element (excluding sex chromosomes) as one asynapsis event and determined the percentage of cells with asynapsis (all data is collected in the linked Dryad Repository).

**Figure 2.**
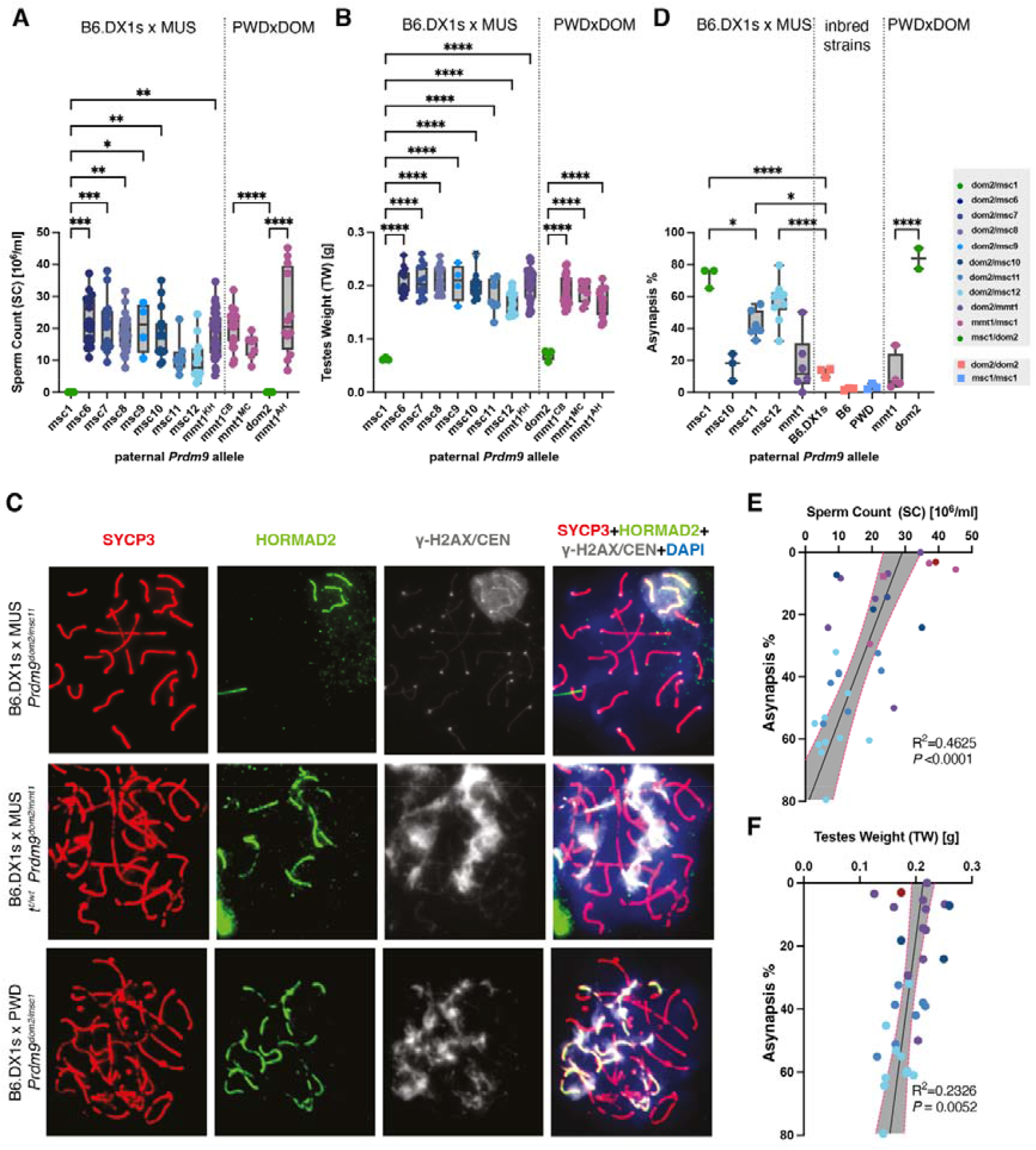
Fertility parameters of intersubspecific F1-hybrids. **(A)** sperm count **(B)** paired testes weights for intersubspecific offspring of (B6.DX1s x wild *MUS*) or (PWD x wild *DOM)* crosses grouped by *Prdm9* genotype and compared to offspring of known hybrid sterility crosses (B6.DX1s x PWD) and (PWD x B6). The asterisks refer to significance values of pairwise ANOVA with Bonferroni correction, with *P<*0.0332 (*), *P<*0.0021(**), *P<*0.0002(***), and *P<*0.0001(****). All hybrid males carry the *Hstx1*^*PWD*^ allele on Chr X. The defects in chromosome asynapsis were assessed by antibody staining for HORMAD2 protein (green), which marks asynapsed autosomal chromosomes in addition to the nonhomologous parts of XY sex chromosomes that are physiologically observed in normally progressing meiocytes. DNA is counterstained with DAPI (blue). Synaptonemal complex assembly was detected via SYCP3 protein immunostaining (red) and yH2AX (grey). At the zygotene/pachytene transition, clouds of yH2AX mark chromatin associated with asynapsed axes (Burgoyne *et al*. 2009; Elinati *et al*. 2017). In addition, localized grey dots represent CEN-labeled centromeres. The panels show spermatocyte spreads of two intersubspecific B6.DX1s x wild MUS, with *Prdm9* genotypes (top) *dom2/msc11* (middle) *dom2/mmt1* and *t-*haplotype, (bottom) *dom2/msc1* of a control F_1_-hybrid male of cross B6.DX1s x PWD. **(D)** Asynapsis phenotypes of intersubspecific F_1_-hybrids grouped by *Prdm9* genotype and parental strains B6DX1s, B6, and PWD **(E)** sperm count and **(F)** paired testes weights correlated with the percentage of asynaptic cells on the Y-axis, red dotted lines represent 95% confidence intervals, ***P*** values, and Pearson values **r** are given on the bottom right

The F_1_ hybrid males with wild MUS alleles *msc11* and *msc12* had a large proportion of asynaptic cells among the total number of evaluated pachynemas (*msc11*, 42.4 ± 8.0 % and *msc12* 57.2 ± 12.5%), these frequencies are significantly elevated, over those in control mice of pure parental subspecies (B6.DX1s and PWD) (as shown in **Figure 2C**). In contrast, hybrids with wild mouse allele *msc10* had lower asynapsis, averaging 16.6 ± 8.6% of cells with asynapsis, not significantly different from fertile controls and significantly lower than sterile controls (**Figure 2C**). Similarly, mice with *t-*haplotypes also show low asynapsis rates (DOM *mmt1* 11.5 ± 12%, MUS *mmt1* 17.4 ± 18.0%). We also tested whether fertility parameters were correlated across all genotypes and indeed observed a weak but significant correlation between high asynapsis with both low sperm count (Pearson R^2^ = 0.46, *P <* 0.0001) (**Figure 2E**) and paired testes weights (Pearson R^2^ = 0.23, *P=* 0.0052) (**Figure 2F**). We also tested the impact of the human variant, which is rescuing sterility when combined with the *msc1’* sterility’ allele (Davies *et al*. 2016), and increased fertility in combination with wild mouse alleles (Mukaj *et al*. 2020). However, the humanized allele did not significantly affect sperm counts, increased paired testes’ weights in a few of the *Prdm9* alleles, but reduced testes weights in mice with *mmt1* on MUS background (**Figure S8**), thus revealing no clear pattern. In summary, while the tested *Prdm9* allelic combinations were either completely fertile or showed a small reduction of fertility, hybrids with *msc11* and *msc12* alleles had the lowest sperm counts and paired testes weights and also showed the highest levels of chromosomal asynapsis. We can therefore conclude that *Prdm9* likely drives this effect, but that any *Prdm9*-dependent effect is either not binomial or affected by modifiers.

### The predominance of *Prdm9* genotype over genetic background

To assess whether potential modifiers on the wild genetic backgrounds affect fertility parameters, we tested whether fertility phenotypes would segregate with the *Prdm9* genotype. In particular, we designed crosses in which the progeny would segregate the PWD *msc1”* sterility” allele and the tested “fertility” alleles on mixed genetic backgrounds, keeping the *Hstx2*^*PWD*^ allele controlled. To do so, we prepared test crosses of either (B6 x DOM) intraspecific hybrid males, which we crossed to PWD females to test the *mmt1* allele on the DOM background. To test MUS alleles, we crossed B6.DX1s females with intraspecific (PWD x MUS) F_1_ hybrid males, whose sperm counts were not significantly different from those of intersubspecific mice (**Figure S8**). In reciprocal orientation (PWD x wild MUS), we crossed F_1_ hybrid females with B6 males. Along the female germline, the *mmt1* allele showed even transmission, equally characteristic of t-haplotype transmission distortion (Lyon 2003). Yet, here, not only *Prdm9* but also X-chromosomes segregate into wild MUS and PWD X-chromosomal haplotypes, of which only the latter carries the *Hstx2*^PWD^ locus (Bhattacharyya *et al*. 2014). To exclusively test the effect of *Prdm9* on wild genomic backgrounds, we only included males with X-chromosomal microsatellite markers SR51, SX69084, and SX65100 alleles indicating the refined *Hstx2*^PWD^ locus (Lustyk *et al*. 2019)(see also **Figure S9**).

Siblings that inherited *Prdm9* alleles *msc6, msc7, msc8*, or the *mmt1* from MUS or DOM all displayed fertility phenotypes within the physiologically normal range, while their brothers with allelic combination *msc1/dom2* were sterile (**Figure 3**), with testes weight and sperm count comparable to laboratory inbred males with the same allelic combination. This shows that hybrid sterility occurs when the allelic combination of the laboratory model of hybrids sterility is inherited, but even when the background is partially wild. As a heterogeneous wild genomic background does not prevent sterility *per-se*, indicating that the effects of high fertility in wild mice hybrids are not primarily due to background effects. Therefore, as genomic modifiers prevent *Prdm9*-dependent hybrid sterility of the *msc1/dom2* allelic combination, hybrid sterility is under oligogenic control, with *Prdm9* as the leading player. Notably, even subtle changes in the *Prdm9* coding sequence produce dramatic effects. This is particularly evident for the *msc6* allele, which differs from the *msc1* allele in only two nucleotides coding for amino acids of the eleventh zinc finger responsible for DNA binding specificity (Figure S4); a slight difference that is nevertheless sufficient in determining whether F1-hybrids are sterile or fertile on the same genetic background. Another example of how small changes can affect fertility can be observed for *msc11*, which displays lower fertility, but equally differs only in a single ZNF from the *Cst* allele, which has been well-characterized in B6xCAST intersubspecific hybrids, and no fertility reduction has been reported (Parvanov *et al*. 2009; Baker *et al*. 2015; Diagouraga *et al*. 2018; Grey *et al*. 2018; Hinch *et al*. 2019; Valiskova *et al*. 2022).

**Figure 3.**
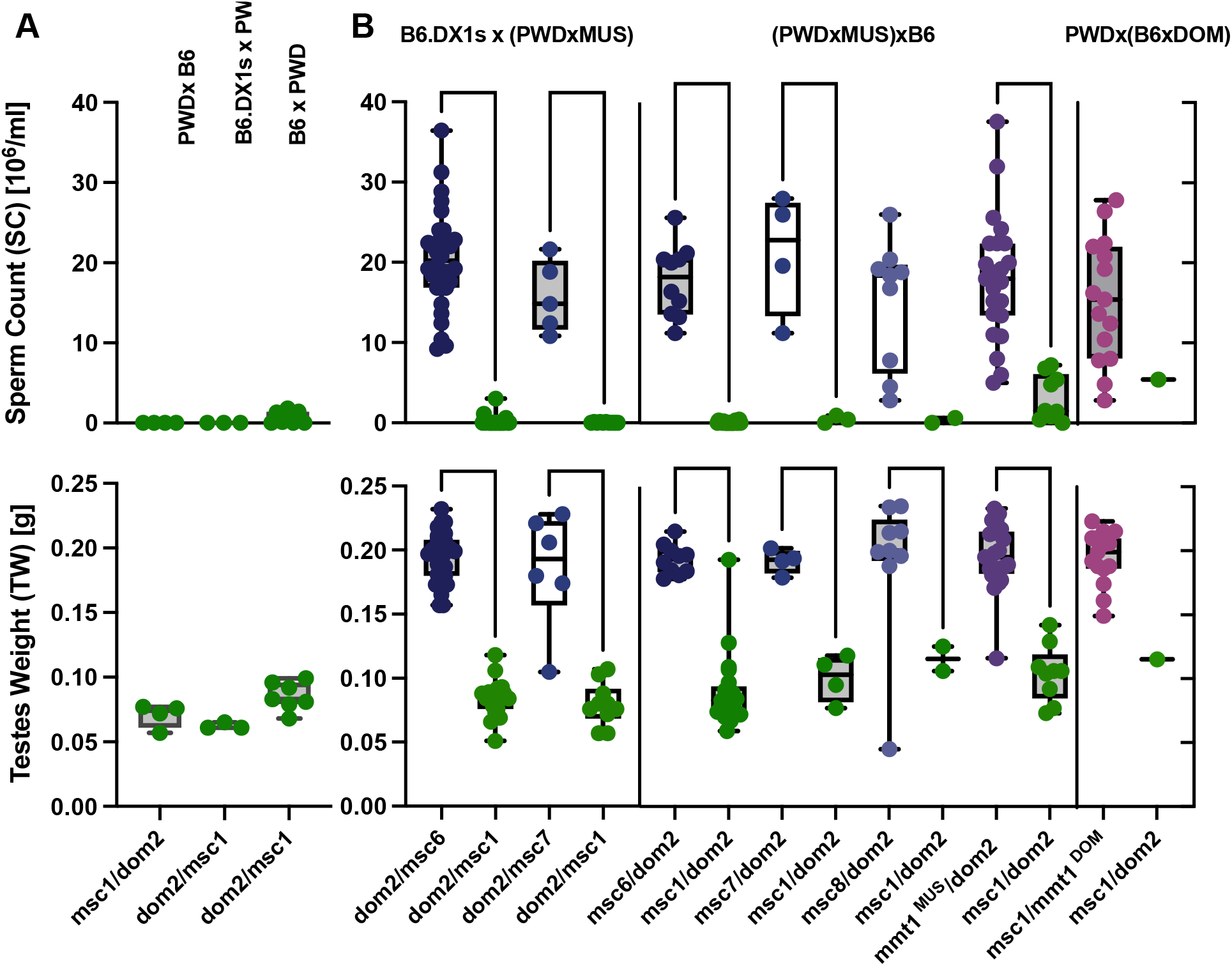
Fertility phenotypes segregate with parental *Prdm9* alleles in reciprocal intersubspecific hybrids. (A) Fertility parameters of control cross with the *Prdm9* allelic combination *dom2/msc1* and Hstx2 allele from PWD or B6. (B) Intersubspecific F_1_ male offspring of **(left)** B6.DX1s females crossed to intrasubspecific MUS males **(middle)**, Intraspecific MUS hybrid females crossed to B6 males **(right)** PWD females crossed to intrasubspecific DOM males. Data were pooled from parents with the same *Prdm9* genotype. Color and statistics, as in Figure 2.

### Variation in X-chromosomal haplotypes in *Prdm9-mediated* sterility

However, even with the allelic combination *msc1/dom2*, not all F_1_ hybrid males are fully sterile. We see a few semi-sterile F1-hybrids with counts of up to 0.2×10^6^ sperm cells. In F_1_ hybrids of laboratory-inbred mice, residual sperm had previously been observed for the same *Prdm9* allelic combination if hybrids had inherited the *Hstx2*^*B6*^ allele (Bhattacharyya *et al*. 2014).

We had assumed a low probability of recombination events in the *Hstx2* locus, as the interval 65-69 Mb on Chromosome X containing the *Hstx2* locus appears as a recombination cold spot (Brick *et al*. 2018; Lustyk *et al*. 2019). However, as masked recombination events are, in theory, possible in intrasubspecific mothers, we inquired into wild X-chromosomal haplotypes of the Kazakh (KH) MUS population. Based on only three microsatellite markers, we identified four distinct MUS X-chromosomal haplotypes: KHa, *KHb, KHc*, and *KHd* (see **Figure S9**). Indeed, we see semi-sterile offspring with an apparent *Hstx2*^*PWD*^ haplotype only in mice where X-chromosomal haplotypes share at least one marker, indicating that masked recombination events within the *Hstx2* locus could have occurred. Therefore, we tested whether wild X-chromosomal haplotypes affected fertility by repeating the *Prdm9 segregation* analyses in males possessing wild X-chromosomal haplotypes. Indeed, just as previously observed with the *Hstx2*^*PWD*^ sterility allele, fertility segregated predominantly by *Prdm9* genotype and irrespective of which X-chromosomal haplotypes the offspring possessed (**Figure S10A-C**). Regardless of X-chromosomal haplotypes, all F_1_-hybrids that inherited the wild MUS allele from their mothers were fertile, while siblings inheriting the maternal *msc1* alleles can be considered infertile. This data shows the absence of genetic modifiers on the wild genetic background also along the maternal germline, but also that the *Prdm9* mediated sterility is not strongly affected by wild X-chromosomal haplotypes. Because comparing all offspring that inherited the *Prdm9* genotype *msc1/dom2*, confirms that whether possessing the *Hstx2*^*PWD*^ X-chromosomal haplotype or any of the tested wild X-chromosomal haplotypes does not significantly affect fertility parameters (**Figure S10D**). Regardless of the X-chromosomal haplotype, not all F1-hybrid males who had inherited the *msc1/dom2 allele* combination were completely sterile, with some nevertheless producing traces of sperm. Notably, these males all had a mother with a t-haplotype. Regardless of X-chromosomal haplotype, offspring with *Prdm9* genotype *msc1/dom2*, whose mothers were t-haplotype carriers, differ significantly in both sperm count and testes weights from mice with the same genotypes whose mothers did not possess t-haplotypes (Mann-Whitney test; SC *P* > 0.0001, TW *P* = 0.0094) (**Figure S10E**). This observation could indicate the presence of *Prdm9* fertility modifier(s) in *t*-carrying populations or, more likely, a *trans*-effect of the *t*-haplotype on homologous Chr 17 in females, analogous to a ‘sperm killer’ effect that causes their transmission ratio distortion in males (Lyon 2003).

### A model for *Prdm9* evolution and hotspot erosion in wild mouse populations

In conclusion, none of the tested novel allelic combinations of wild *Prdm9* alleles produces completely sterile F_1_-hybrid male offspring, fully consistent with the low levels of sterile hybrids reported in the wild (Turner *et al*. 2012). Instead, we see either completely fertile hybrids or a significant reduction of *Prdm9*-dependent fertility and high levels of asynapsis in intersubspecific hybrids. We have observed that possessing a heterogeneous wild genomic background does not appear to prevent sterility *per-se*. The *msc1/dom2* allelic combination alone is necessary and sufficient to create hybrid sterility, confirming that the effects of high fertility in wild mice hybrids are not primarily due to background effects in our crosses. Furthermore, our segregation analyses show that fertility or sterility phenotypes segregated purely with the *Prdm9* genotype, thus fully supporting that hybrid sterility is under oligogenic control with *Prdm9* as the leading player. Taken together with previous data (Mukaj *et al*. 2020), it appears that sterility alleles of *Prdm9* may be rare in natural populations of mice.

It was hypothesized that the molecular mechanism of *Prdm9* action could be related to the evolutionary divergence of homologous genomic sequences in *DOM* and *MUS* subspecies (Davies *et al*. 2016) and, more specifically, to the phenomenon of historical erosion of genomic binding sites of PRDM9 ZNF domains (Baker *et al*. 2015) caused by repeated biased gene conversion. Full sterility was observed in hybrids of inbred mice, where hotspot erosion may be exacerbated. Highly inbred genomes, such as the genomes of the B6 or C3H strain, which show severe or moderate sterility in combination with PWD, are not comparable to heterozygosity levels in initiation motifs in wild mouse genomes. A genome that initially possessed wildtype levels of heterozygosity becomes sequentially homozygous through inbreeding, and this process should equally affect PRDM9 binding sites. Therefore, once a strain is fully inbred, there should be no more heterozygous binding motifs, and erosion should cease completely or at least slow down substantially, as it is now based only on rare mutations with stochastic placement within binding sites. Therefore, substantial erosion of binding sites must have occurred before inbreeding. Given the high frequency at which they are found in modern inbred strains, these alleles should have existed for long evolutionary timescales and at larger population frequencies before the strains were established by inbreeding. The *msc1* sterility allele from PWD/Ph is also found in the STUS/Jpia strain, and the *dom2* sterility allele is found in several laboratory inbred strains, including C57Bl6/J, A/J, DBA/2J, LP/J, 129s1/SvlmJ, C57BL/10J and C57BR/J and sterility allele *dom3* is not only found in the C3H, WSB/EiJ, PERA/EiJ, SK/CamEiJ, CBA/J, BDP/J, P/J, SJL/J, and RFM/J laboratory inbred strains but also in the wild-caught SIN/Jpia, SIT/Jpia, and SCHEST/Jpia strains (Mukaj 2020).

Allele *msc11*, which displayed the second-lowest sperm count and second-highest asynapsis in this study, is among the alleles with the least evolutionary divergence from the last common ancestor. The *msc11* allele was previously identified as 7mus1 in MUS from Austria and is highly similar to CAS alleles, differing by only single amino-acid substitutions from the *Cst* allele and a wild mouse allele from Iran. The similarity of *msc11* to alleles found in the CAS subspecies may indicate that this allele may have originated in CAS, which is phylogenetically older than MUS, and that it may have already been present before the subspecies split. Furthermore, msc11 appears to have had a large distribution range, which, taken together with older phylogenetic age, would predict a high level of hotspot erosion. However, these observations are equally true for the *msc10* allele, which only showed a low level of Asynapsis in this study. Furthermore, the t-haplotype associated *mmt1* allele, which can be considered an intraspecies *Mus musculus* allele rather than a MUS or DOM allele, is one of the evolutionarily oldest alleles and globally occurs at high population frequencies. However, as it is an introgression allele, not its evolutionary age but the time of introgression is relevant for when it started eroding binding sites in the genomes in question. Given that it is typically present at high population frequencies, large-scale erosion of binding sites should have occurred. However, given that the same allele is present in multiple subspecies, symmetric erosion would occur on both genomes simultaneously, which may nevertheless ensure symmetric hotspot binding, despite leading to a lower number of hotspots overall. Finally, the homozygous lethality of t-haplotype carriers ensures that there is always *Prdm9* heterozygosity. As long as the second allele possesses sufficient binding sites, this heterozygosity alone may be sufficient to rescue fertility in t-haplotype carriers.

To avoid negative selection of a complete loss of recombination hotspots over time, *Prdm9* evolves rapidly with protein variants behaving like the predator and specific motifs as prey following Red Queen dynamics (Latrille *et al*. 2017). *Prdm9* evolves rapidly, driven by *de-novo* recombination events during meiosis, including crossover and gene-conversion-like events (Jeffreys *et al*. 2013). Given the rapid evolution of *Prdm9*, substantial hotspot erosion may not occur in any given population if large numbers of variants are present at low population frequencies. And indeed, LD recombination maps on the source populations of mice from this study reveal only weak conservation of recombination maps at the broad and the fine scale, even at the population level, with most hotspots unique to each population (Wooldridge and Dumont 2022). In this scenario, population-specific erosion of hotspots would be expected, leading to variation in the fertility of the same *Prdm9*-allele in different populations. Suppose Individual hotspots specific to only one variant would experience only a low historical erosion level within a given population. In that case, only alleles, which have been present for long evolutionary timescales, have large distribution ranges, and are present at high population frequencies are expected to also have eroded recombination landscapes and show reduced fertility. Variation between fertility parameters of the same allele on different backgrounds has indeed been documented for both the *dom3* allele and *msc5* allele before (Mukaj *et al*. 2020).

Furthermore, while the *msc11* allele appears to be evolutionarily old, our phylogenetic analyses revealed several previously identified hybrid sterility alleles such as *msc5* from BULS, BUSNA, and STUF and *msc1* from STUS and PWD (Mukaj *et al*. 2020), not to be among those alleles that were most closely related to a common ancestor. The *msc1* allele from PWD and the *msc2* allele found in the SKE/Jpia strain (Mukaj *et al*. 2020) share a node within a phylogenetic branch exclusively containing MUS alleles, while *msc5* is located on a distant branch. However, strikingly, *Prdm9* alleles for which full hybrid sterility was observed to date code for a long stretch of five identical ZNFs. The *msc12* allele, which had the lowest sperm count and the highest level of chromosomal asynapses in this study, is identical in four of the five ZNFs and differs only in a single amino-acid residue in the fifths. At least in human hotspots, many similar variants were able to activate the same recombination hotspot motif (Berg *et al*. 2010; Berg *et al*. 2011). Therefore, diverse hybrid sterility alleles may lead to the activation, and subsequent erosion of the same DNA initiation motif, particularly in scenarios where not all ZNFs are equally important in DNA binding. Indeed, our data also indicate that particular zinc fingers appear more critical in DNA-binding or in altering binding affinity than others. Related *Prdm9* alleles differing by two single-nucleotide polymorphisms display dramatically different fertility phenotypes. While the *msc6* allele from the MUS population from Kazakhstan differed only in a single ZNF from the *msc1* allele, siblings inheriting the *msc6* allele are fertile, while those inheriting the *msc1* are sterile, making differences between ZNF11 solely responsible for the fertility difference, on the same genetic background. A similarly slight divergence between ZNF12, coded for by the *msc4* allele from the SKE/Jpia stain and the *msc1* allele from PWD, did not alter sterility phenotypes.

In conclusion, our data are consistent with an increasing fertility decline with gradual hotspot erosion over time instead of a binomial effect where mice are either fully sterile or fully fertile. It appears that hotspot erosion can occur either for evolutionarily old alleles with a small evolutionary divergence from the last common ancestor or alleles with significant motif overlap across populations. The rapid evolution of Prdm9 could therefore explain the lack of hybrid sterility phenotypes in wild mice. Given the high natural divergence of *Prdm9* in wild mouse populations, substantial hotspot erosion may not occur in any given population if large numbers of variants are present at low population frequencies. In this scenario, wild mouse genomes may only rarely encounter the same PRDM9 variant for several generations, and most individual hotspots specific to only one variant would experience a low historical erosion level. However, in speciation scenarios with significant bottlenecks, reduced *Prdm9* diversity within a small population size would predict extensive hotspot erosion over fewer generations, leading to hybrid sterility phenotypes upon secondary contact.

## Availability of data and materials

Nucleotide sequences of *Prdm9* alleles were deposited to Genbank under accession numbers OQ055171 - OQ055188. The fertility datasets generated and analyzed during the current study are available as a Dryad repository under DOI 10.5061/dryad.bzkh189cm. The R package to calculate the genetic distance between complex repeats is available at https://gitlab.gwdg.de/mpievolbio-it/repeatr.

## Acknowledgments

We would like to thank Peter Donnely for the transgene *Prdm9*^*tm1*.*1*^*(PRDM9)*^*Wthg*^ (“humanized”) strain and Attila Toth for the HORMAD2 antibody. We would like to thank Christine Pfeifle and the entire mouse house team at the MPI in Plön for their help with the mouse breeding and maintenance. We are grateful to Heike Harre for fertility phenotyping, Vladana Fotopulosova for help with Cytology, and Nicole Thomsen for DNA extraction and genotyping.

## Author contributions

KFNA, ED, KKU, and AM acquired, analyzed, and interpreted data. JF, EP, and LOH conceived the project, designed the work, supervised data acquisition, and analyzed and interpreted the data. LOH wrote the manuscript. All authors have read and approved the final manuscript.

## Supplementary Data

**Table S1:**
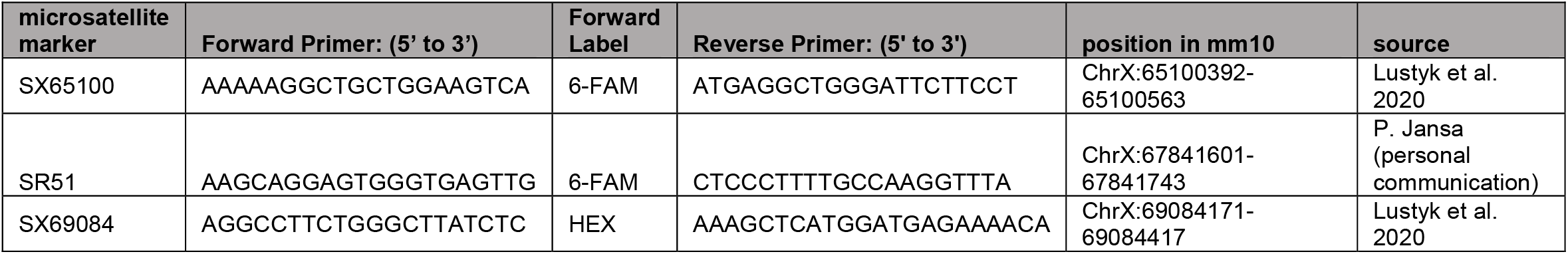
Primers used to genotype the *Hstx2* interval.

**Figure S1:**
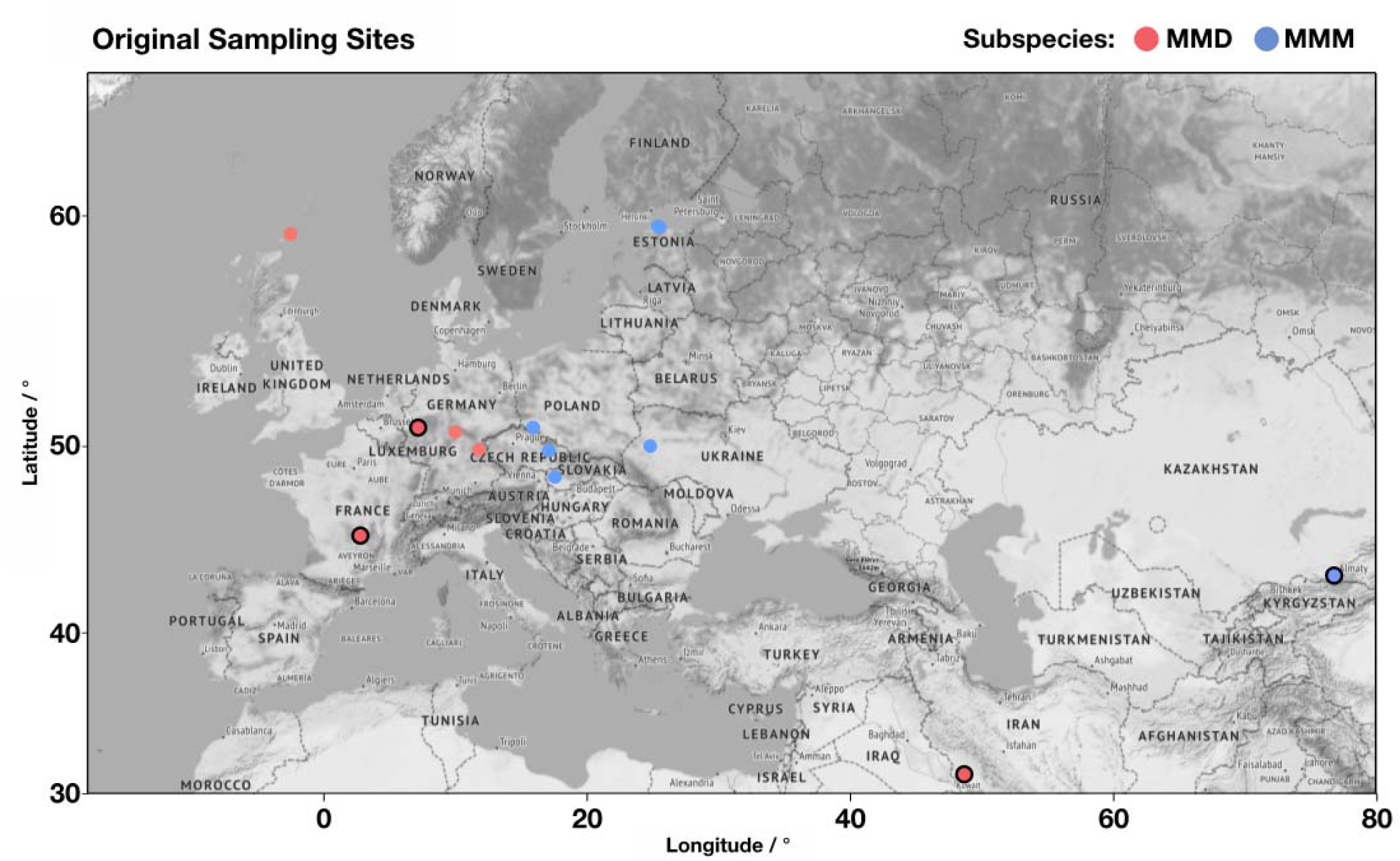
Distribution of original sampling sites of MUS (blue) and DOM (red), with framed circles representing mice evaluated for hybrid sterility in this study, and plain circles from (Mukaj *et al*. 2020)

**Figure S2:**
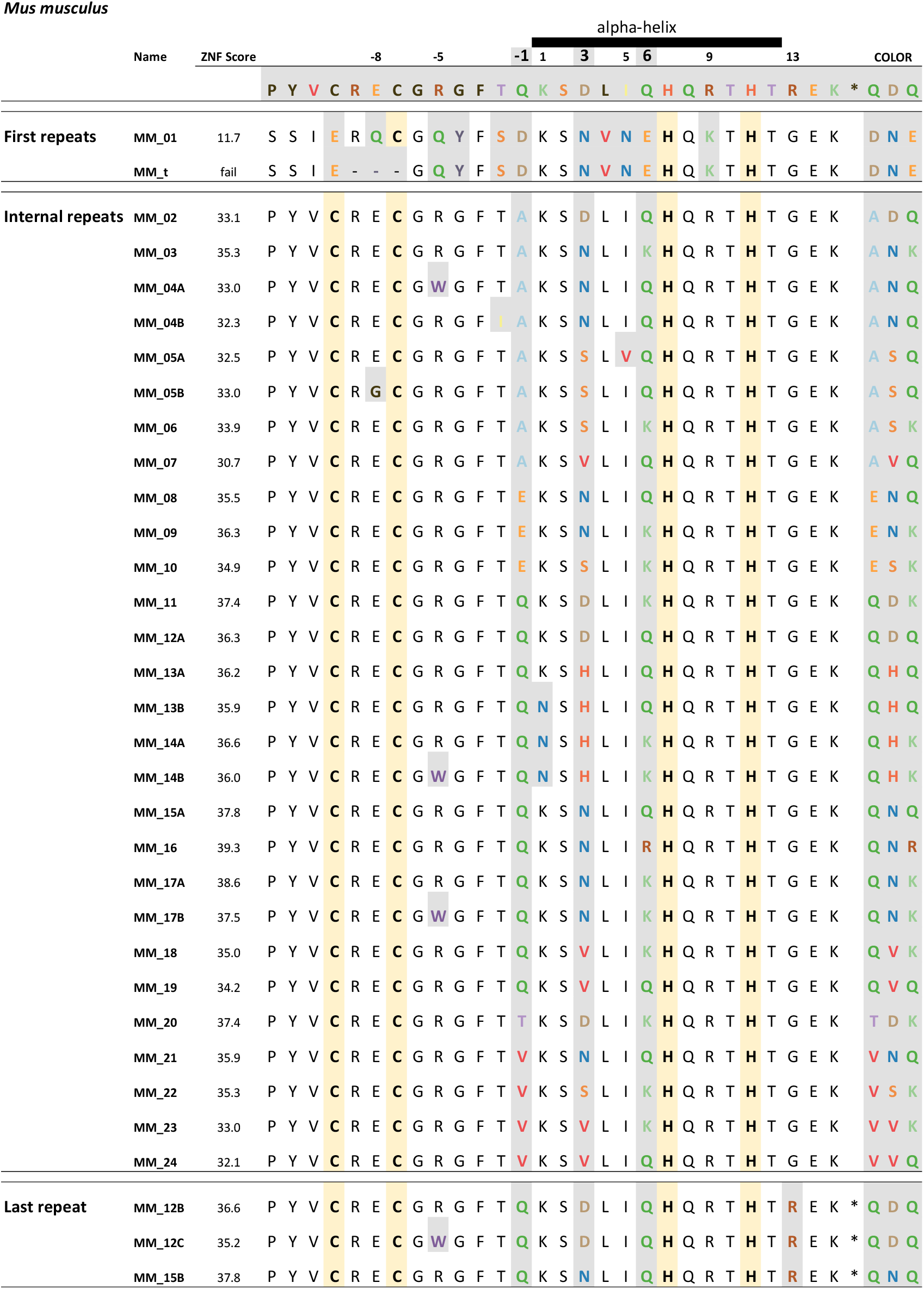
Types of C_2_H_2_ zinc fingers found in PRDM9 MUS and DOM. (from this study and (Mukaj *et al*. 2020). The variable positions (shaded in grey) responsible for the DNA-binding specificity of each Zinc-finger are used as acronyms on the right. Amino acids were colored based on their chemical properties to allow visual distinction of zinc-fingers.

**Figure S3:**
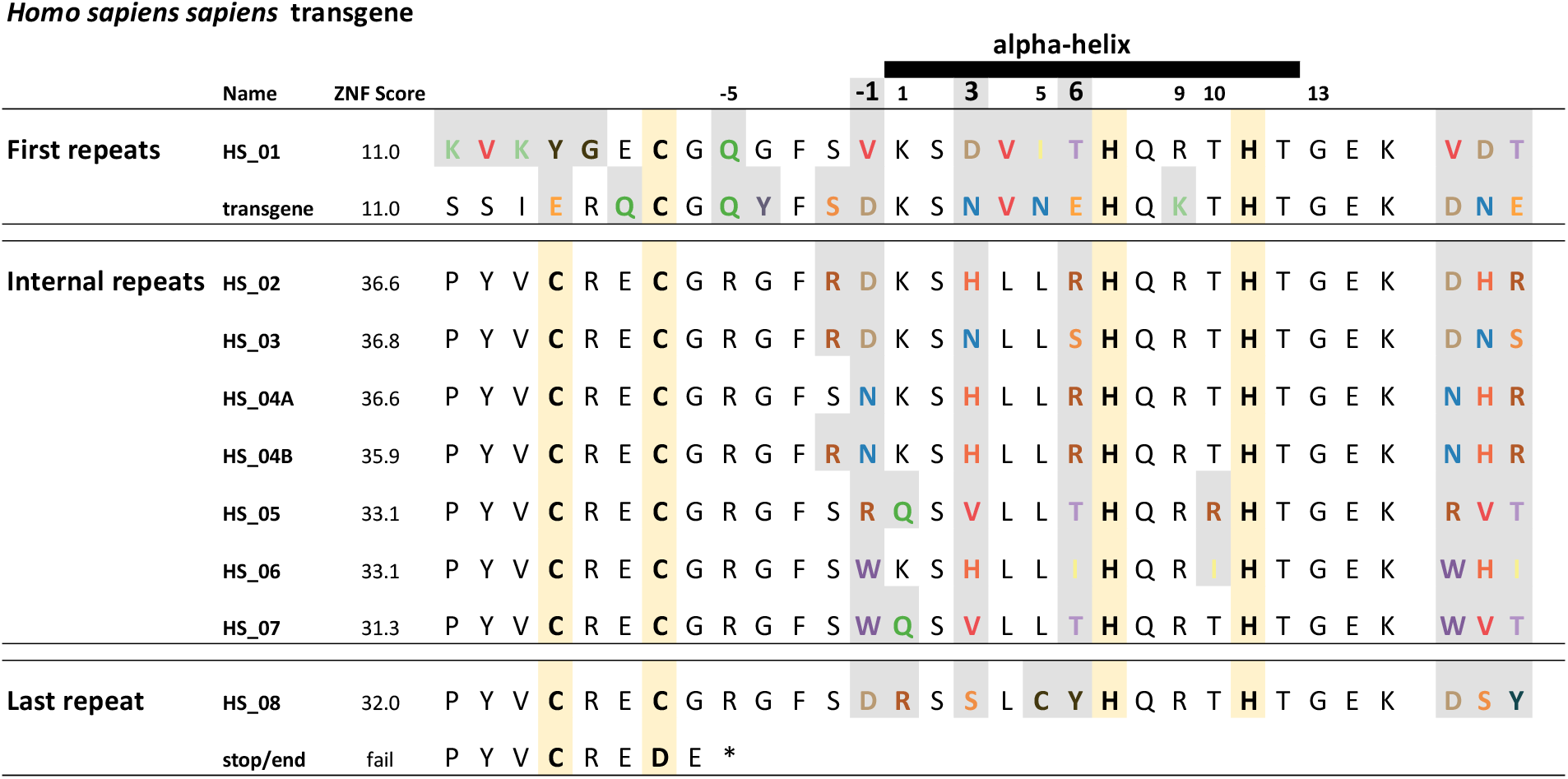
Types of C_2_H_2_ zinc fingers found in “humanized” PRDM9 variant “B.” The variable positions responsible for DNA-binding specificity (shaded in grey) are shaded based on the chemical properties of the amino acid in this position and summarized and used as acronyms on the right.

**Figure S4:**
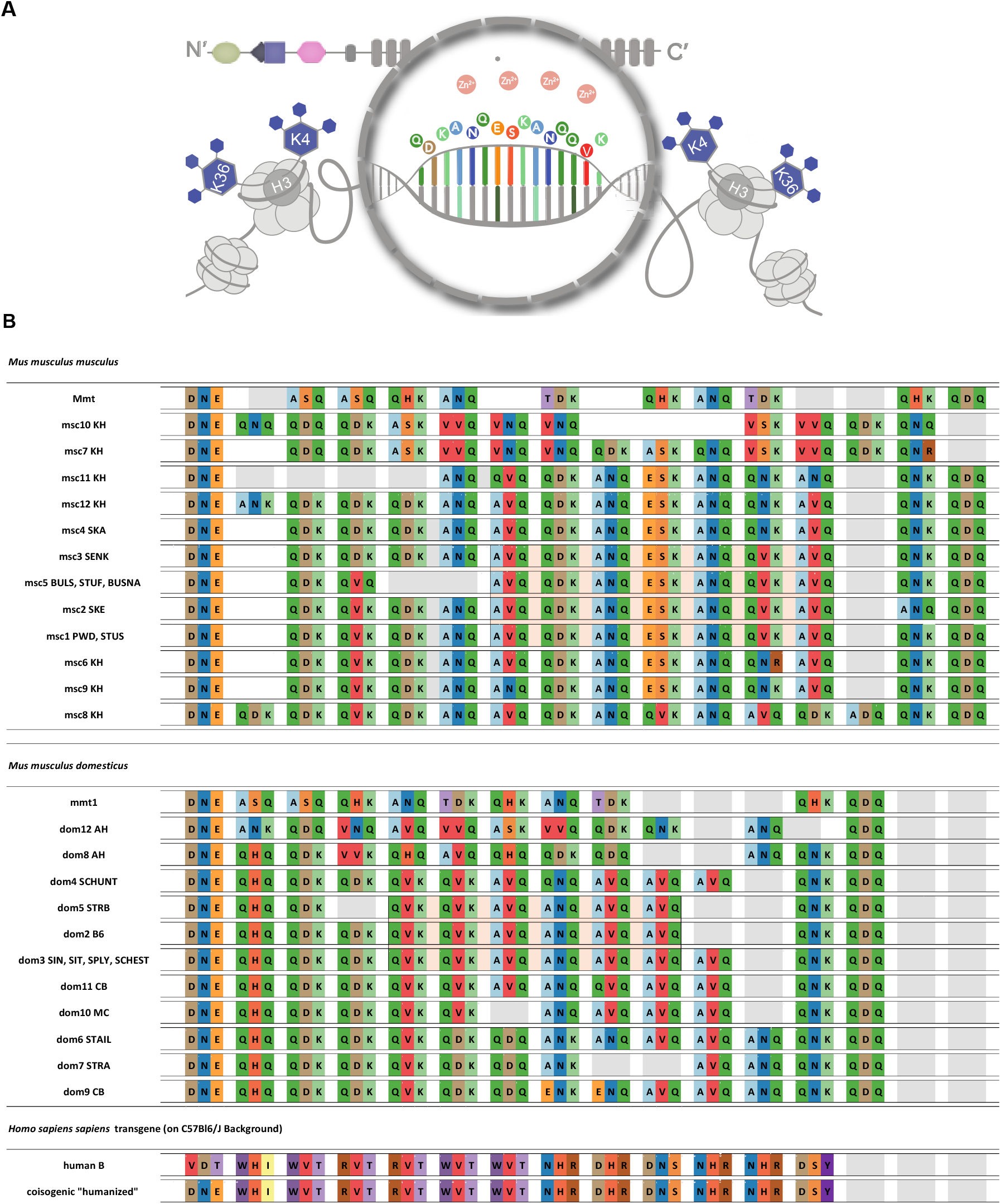
Types of C_2_H_2_ zinc finger arrays studies for hybrid sterility phenotypes (A) principle of PRDM9 binding to target DNA, contact residues of C_2_H_2_ zinc fingers are in positions −1, 3, and 6 of the alpha-helix. **(B)** Representation of all C_2_H_2_ zinc finger arrays using only acronyms of the amino-acid positions responsible for DNA binding (as in **Figure S2** and **Figure S3 Error! Reference source not found**.)

**Figure S5:**
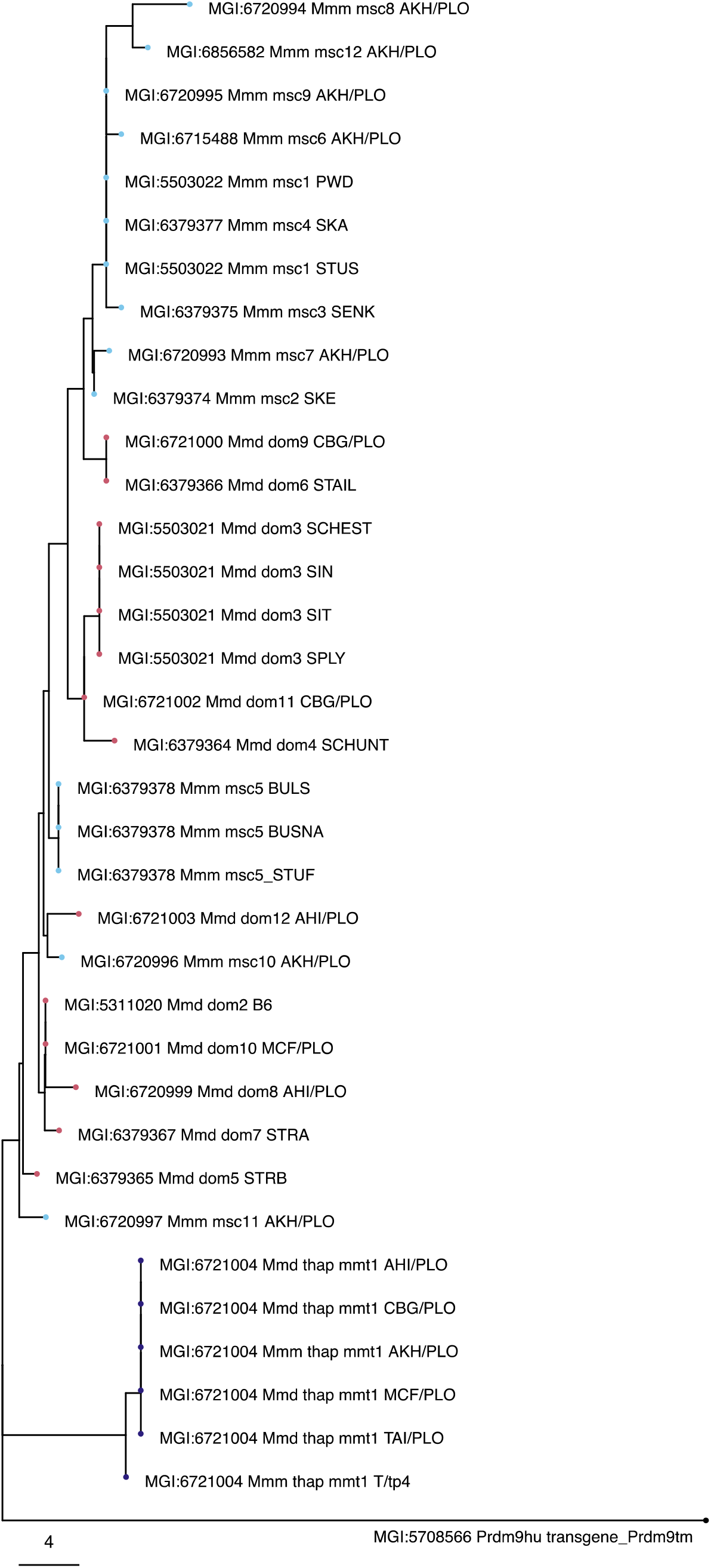
Neighbor-joining *Prdm9* phylogeny based on Hamming. Distances without hypervariable nucleotides that code for the amino acids responsible for DNA binding specificity of PRDM9 and that are known to be under strong positive selection. Data from this study and from (Mukaj *et al*. 2020) were used, red nodes for DOM and blue nodes for MUS, with purple nodes depicting alleles on t-haplotypes found in MUS and DOM.

**Figure S6:**
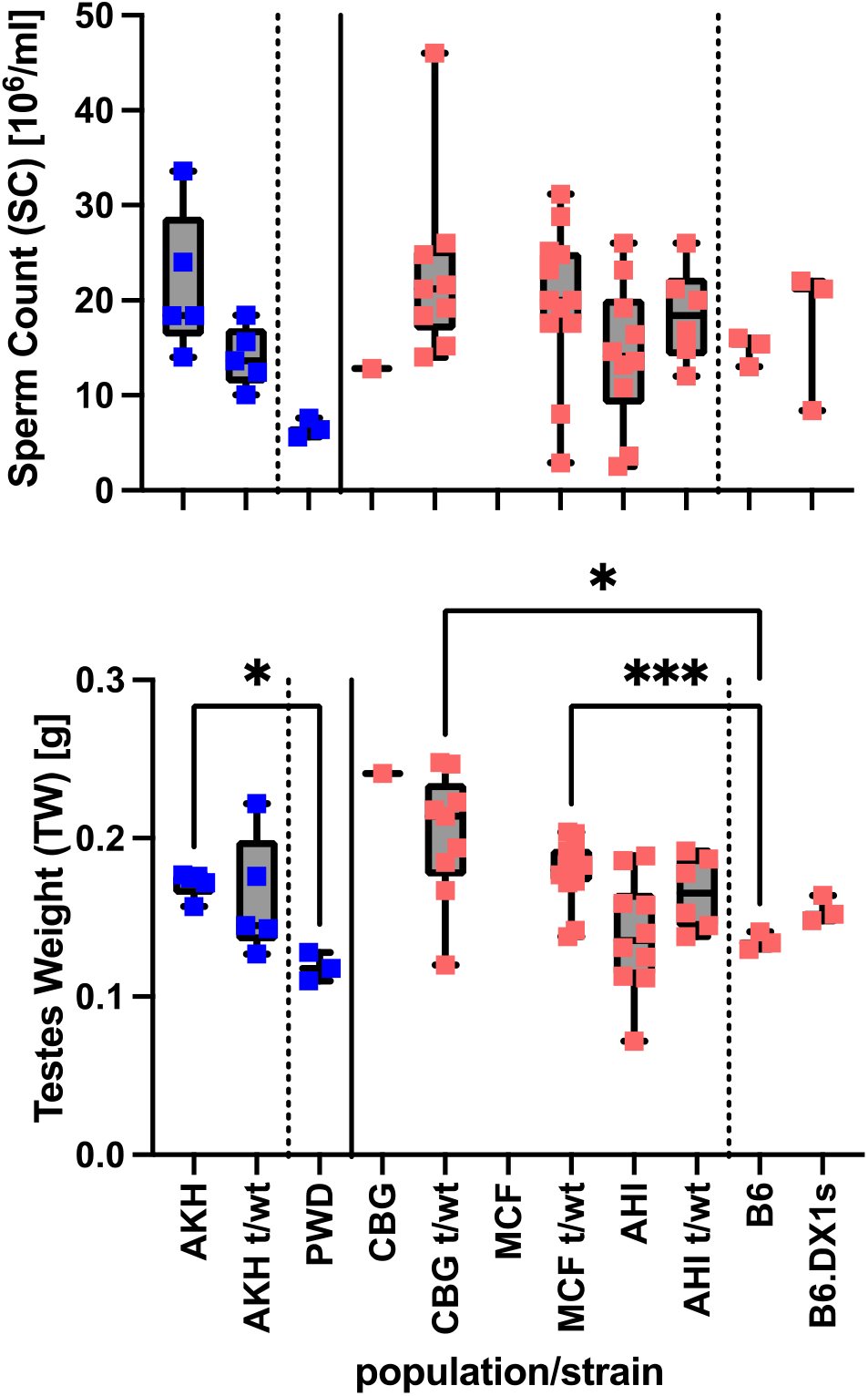
Fertility Parameters of wild mice populations in Ploen, Germany. **MUS**; **AKH** Almaty, Kazakhstan; **DOM**; **AHI** Ahvaz, Iran; **CBG** Cologne-Bonn Germany; **MCF** Massif-Central France; **t/wt** t-haplotype genotype. Statistics as described previously.

**Figure S7:**
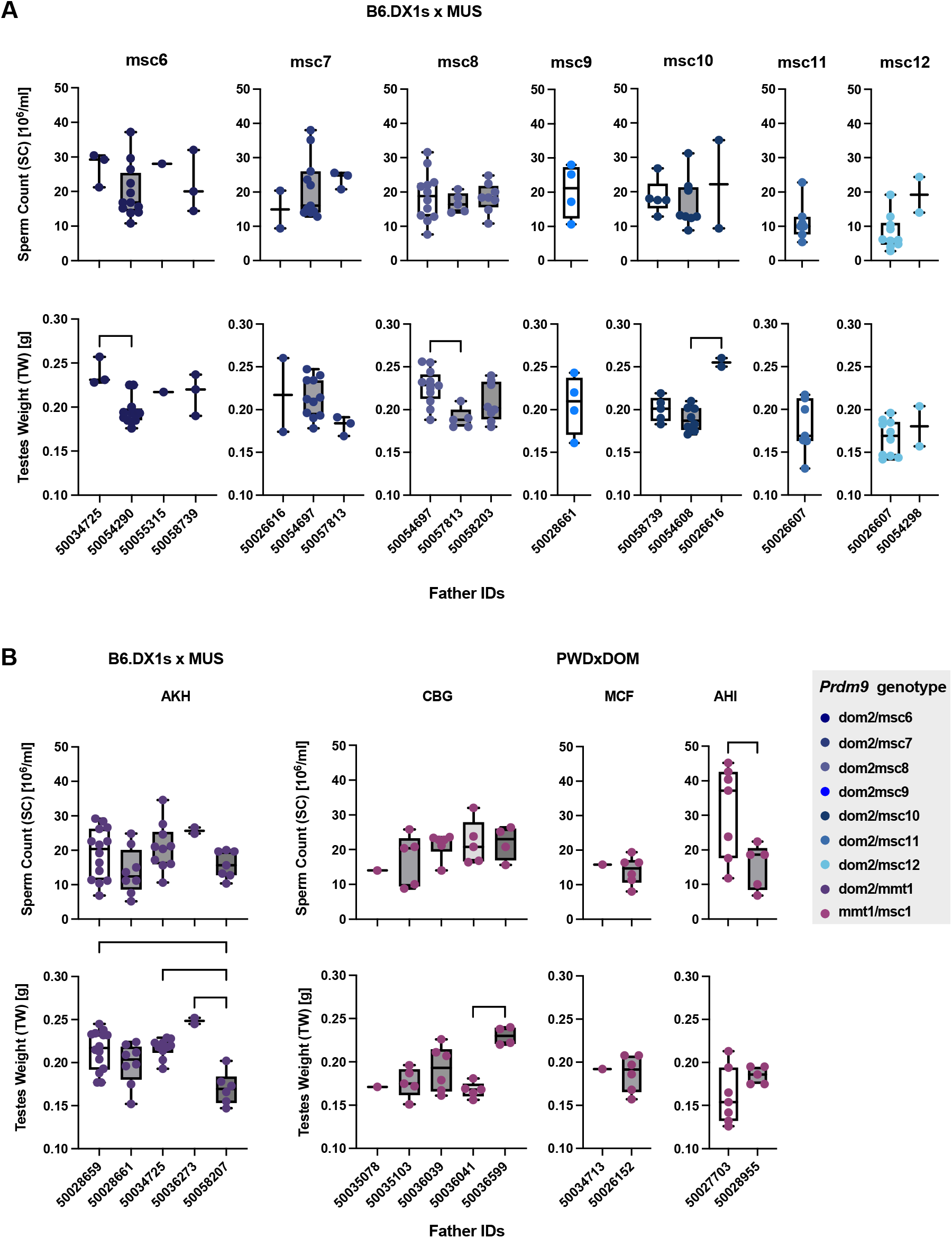
Fertility phenotypes of hybrid offspring grouped by the genotype inherited from the sire in the cross. **(A)** MUS from Kazakhstan without t-haplotypes, all offspring are sorted by Prdm9 genotype, only one sire (50054290) was homozygous for Prdm9, all others heterozygous for different Prdm9 alleles **(B)** Offspring of sires with t-haplotypes, grouped by population origin of the sire (left) MUS from Kazakhstan (right) DOM from Cologne-Bonn, Germany (CBG), Massif-Central, France (MCF) and Ahvaz, Iran (AHI) (right). Data pairs were compared using Welch’s t-test, and multiple comparisons were performed using ANOVA with Kruskal-Wallis test, corrected for multiple comparisons using Dunn’s test. Only significant values are shown on the graph with P<0.0332 (*), P<0.0021(**), P<0.0002(***), P<0.0001(****).

**Figure S8:**
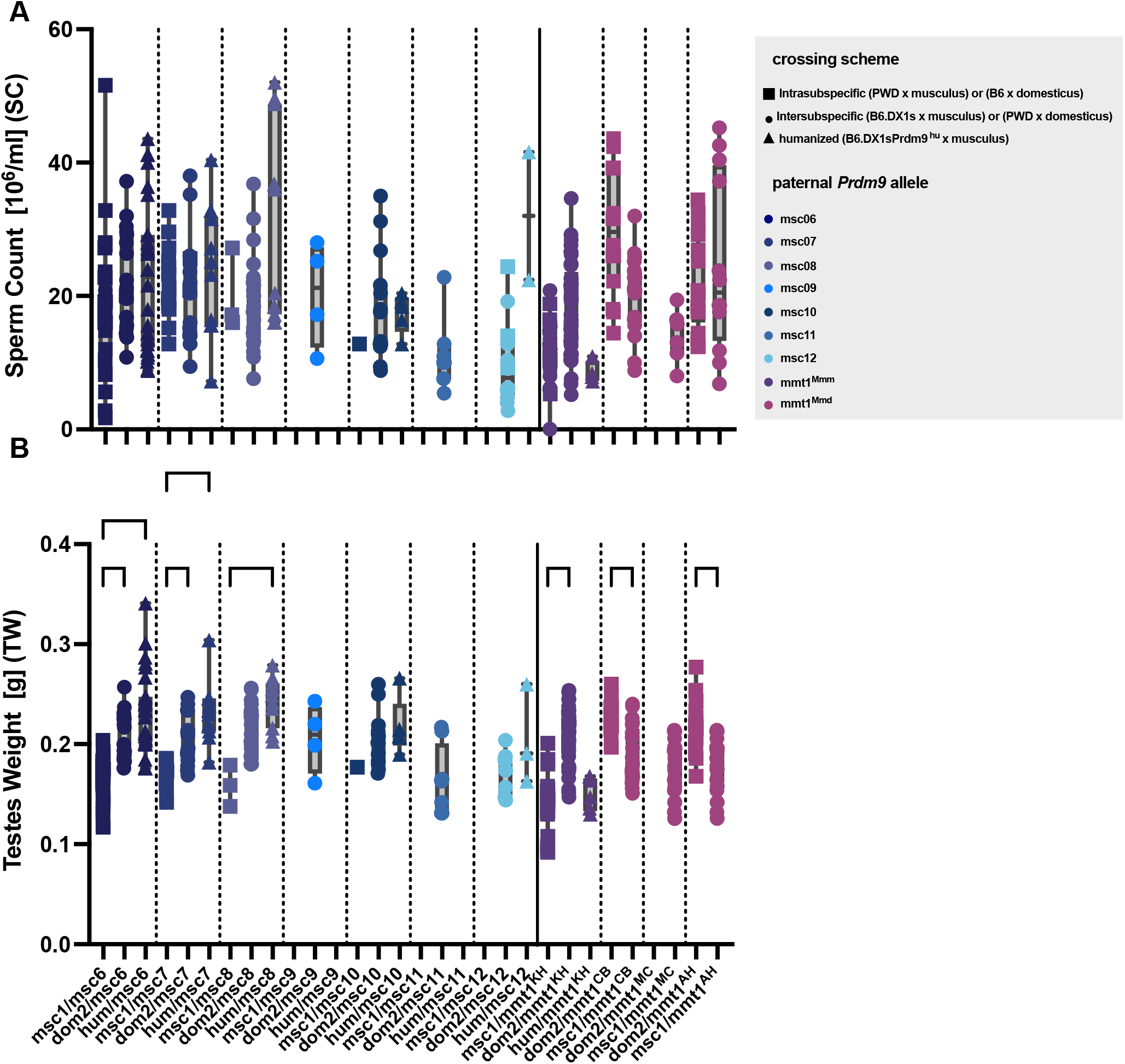
Fertility parameters of intra- and intersubspecific F_1_-hybrids. **(A)** sperm count **(B)** paired testes weight of intersubspecific and Intrasubspecific mice. Paternal Prdm9 alleles are distinguished by colors, while the shape of the data points distinguishes crossing schemes. (Square data points) Intrasubspecific male F_1_ offspring of PWD females crossed to MUS males, or Intrasubspecific B6 females crossed to DOM males. Two types of interspecific crosses were performed; MUS males were crossed to B6.DX1s females and wild DOM males were crossed to PWD females (circles as data points, also shown in **Figure 2**); for the MUS males, a third cross was performed where they were mated with B6.DX1s. Prdm9^hu^ females (triangular data points). Since most males used as sires were heterozygous for different allelic combinations of *Prdm9*, data from several crosses was pooled by *Prdm9* genotype of the F_1_-hybrid offspring, and the significance of differences in fertility phenotypes was evaluated using ordinary one-way ANOVA with Bonferroni correction as before.

**Figure S9:**
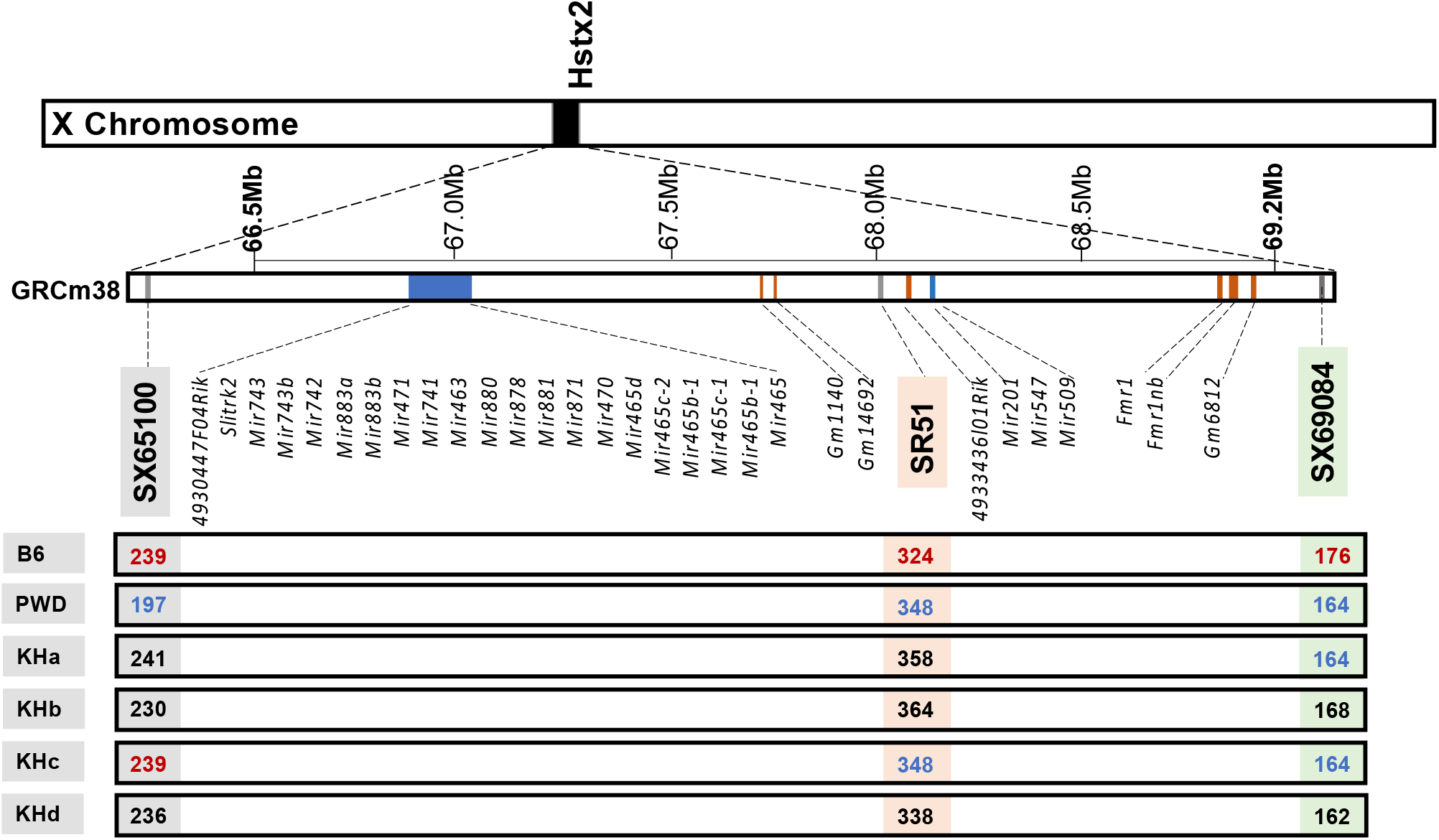
X-chromosomal haplotypes across the refined *Hstx2* locus from (Lustyk et al. 2019) based on allelic variation at microsatellite markers SX65100, SR51, and SX69084 in laboratory and wild mice. All genes and microRNAs annotated within this locus are shown in the mm38 reference genome.

**Figure S10:**
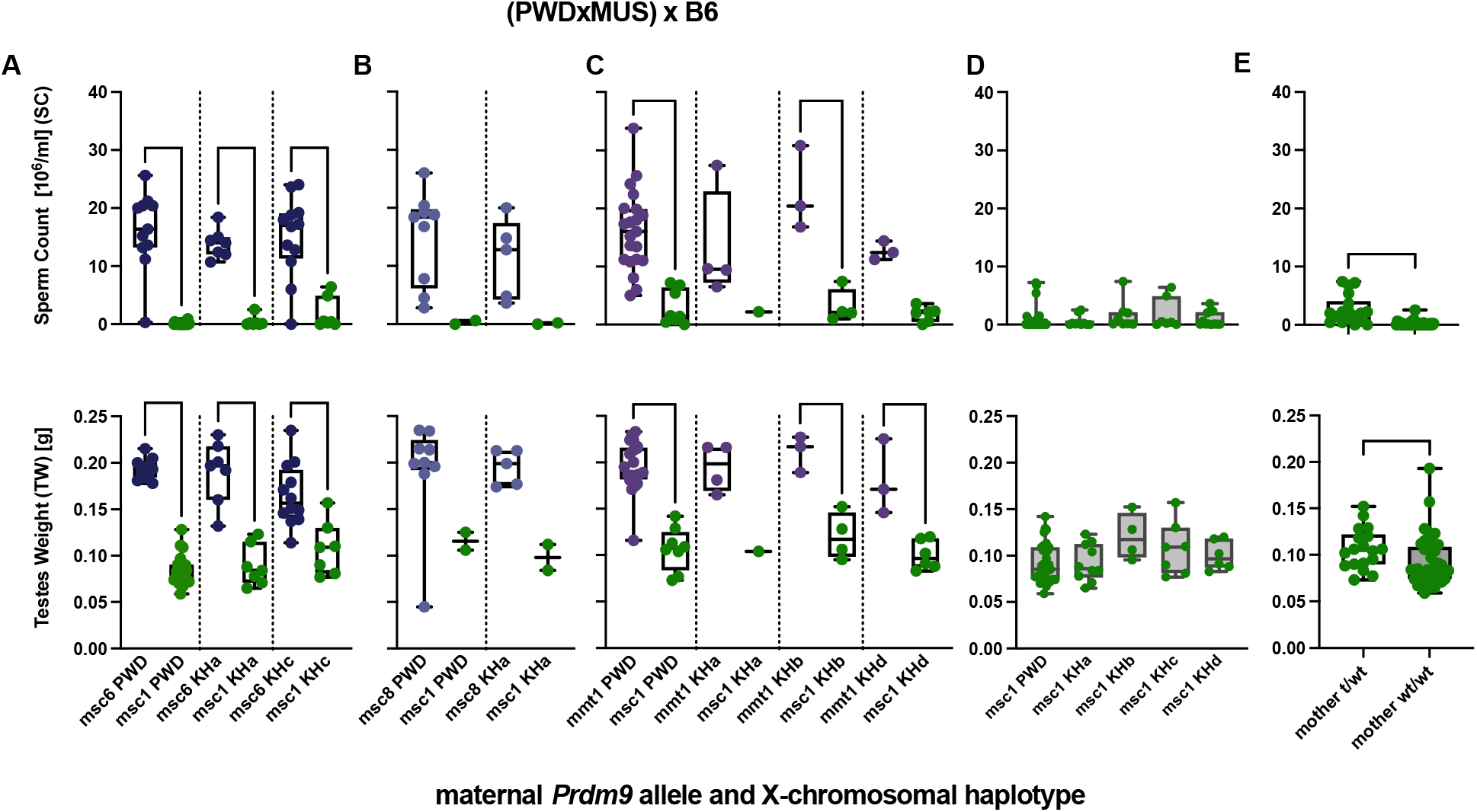
Segregation analyses of maternal *Prdm9* genotype together with the variation of wild X-chromosomal haplotypes. Intersubspecific F_1_ male offspring of Intraspecific MUS hybrid females crossed to B6 males (as in **Figure 3B**) that inherited wild MUS X-chromosomal haplotypes. X-axes depict both the maternal *Prdm9* allele and wild MUS X-chromosomal haplotype, defined as shown in (**Figure S9**). Fertility parameters, sperm count (top), and paired testes weights (bottom) are shown, and the significance of differences in fertility phenotypes was evaluated as before. (A-C) Segregation analyses grouped by maternal *Prdm9* allele combination (A) maternal *msc1* and *msc6* alleles (B) maternal *msc1* and *msc8* alleles (C) *msc1* and *mmt1*^*KH*^ alleles (D-E) Pooled data of hybrids from different mothers that all inherited the same msc1/dom2 allelic combination (D) pooled by X-chromosomal haplotype (E) pooled by whether mothers possessed t-haplotypes.

